# Ancestry-specific polygenic scores and SNP heritability of 25(OH)D in African- and European-ancestry populations

**DOI:** 10.1101/596619

**Authors:** Kathryn E. Hatchell, Qionshi Lu, Scott J. Hebbring, Erin D. Michos, Alexis C. Wood, Corinne D. Engelman

## Abstract

**Context:** Vitamin D inadequacy, assessed by 25-hydroxyvitamin D [25(OH)D], affects around 50% of adults in the United States and is associated with numerous adverse health outcomes. Blood 25(OH)D concentrations are influenced by genetic factors that may determine how much vitamin D intake is required to reach optimal 25(OH)D. Despite large genome-wide association studies (GWASs), only a small portion of the genetic factors contributing to differences in 25(OH)D levels has been discovered.

**Objective:** Therefore, knowledge of a fuller set of genetic factors could be useful for risk prediction of 25(OH)D inadequacy, personalized vitamin D supplementation, and prevention of morbidity and mortality from deficient 25(OH)D.

**Design:** Using PRSice and weights from published African- and European-ancestry GWAS summary statistics, ancestry-specific polygenic scores (PGSs) were created to capture a more complete set of genetic factors.

**Patients or Other Participants:** Participants (European ancestry n=9,569, African ancestry n=2,761) came from three cohort studies.

**Main Outcome Measure(s):** Blood concentrations of 25(OH)D.

**Results:** The PGS for African ancestry was derived using all input SNPs (a p-value cut-off of 1.0) and had an R^2^ of 0.3%; for European ancestry, the optimal PGS used a p-value cut-off of 3.5×10^−4^ in the target/tuning dataset and had an R^2^ of 1.0% in the validation cohort. Those with highest genetic risk had 25(OH)D that was 2.8-3.0 ng/ml lower than those with lowest genetic risk (p=0.0463 to 3.2×10^−13^), requiring an additional 467 to 500 IU of vitamin D intake to maintain equivalent 25(OH)D.

**Conclusions:** PGSs are a powerful predictive tool that could be leveraged for personalized vitamin D supplementation to prevent the negative downstream effects of 25(OH)D inadequacy.

## Introduction

Vitamin D inadequacy, using the Institute of Medicine definition of a 25-hydroxyvitamin D [25(OH)D] concentration less than 20 ng/mL, affects almost 50% of adults in the United States, with higher prevalence in those with darker skin tones (1–3). Observational studies show associations between low vitamin D concentrations and numerous adverse health outcomes, including autoimmune diseases, migraines, hypertension, dyslipidemia, cardiovascular events, and cardiovascular mortality (1,3–9). These studies are supported by recent Mendelian randomization studies which provide evidence for a causal relationship between low vitamin D concentrations and increased risk of obesity, ovarian cancer, hypertension, lower cognitive function during aging, multiple sclerosis, and all cause and cancer mortality (10–16). Furthermore, some clinical trials have shown that vitamin D and calcium supplementation are important in the prevention of fractures and cardiovascular risk factors, while vitamin D supplementation alone may lower risk of cancers, diabetes and depression and may reduce inflammation and improve lung function in patients with cystic fibrosis (7,17–25). Recent results from the Vitamin D and Omrega-3 Trial (VITAL) showed null associations between vitamin D supplementation and cancer or cardiovascular disease. However, study design limits the interpretability of these findings; for example individuals with adequate 25(OH)D concentrations were included, and outside use of vitamin D before and during the trial were not restricted (26). Avoiding vitamin D inadequacy is important, however, as 25(OH)D concentrations over 50 ng/mL have been associated with increased morbidity and mortality (3,27). Clinical trials of vitamin D have shown that individual response to vitamin D supplementation is highly variable (28,29). 25(OH)D concentrations are influenced by genetic factors and genetic variants may determine how much vitamin D intake is required to reach an optimal 25(OH)D blood concentration (30–33). Therefore, knowledge of the genetic determinants of 25(OH)D concentrations could be useful for prediction of risk for vitamin D inadequacy, personalized vitamin D supplementation, and subsequent prevention of vitamin D associated morbidity and mortality due to 25(OH)D deficiency.

Variation in or near twelve genes (*A2BP1, AMDHD1, ANO6/ARID2, CYP2R1, CYP24A1, DAB1, DHCR7, GC, GPR114, HTR2A, KIF4B*, and *SEC23A*) has been associated with serum 25(OH)D at genome-wide levels of significance through published genome-wide association studies (GWASs) in those of European or African ancestry (1,34–37). However, only single nucleotide polymorphisms (SNPs) in or near four of these genes have been replicated (*CYP2R1, CYP24A1, DHCR7*, and *GC*), and together account for a small portion of the variation in 25(OH)D concentrations, about 2.8% compared to the estimated 20-40% heritability (1,31,36). Such “missing heritability” is common in complex traits, and could, in part, be attributed to many SNPs with small effects that do not reach a stringent genome-wide significance threshold (38). A polygenic score (PGS), by comprising the weighted sum of trait-associated alleles, may capture more trait variation than individual SNPs alone. PGSs have been shown to be more powerful than individual SNP-based testing, are used in a wide variety of statistical techniques (e.g., Mendelian randomization), and have shown clinical promise, predicting Alzheimer’s disease incidence before the onset of symptoms that would result in a clinical diagnosis, and for dosing of antifibrinolytic drugs based on activated partial thromboplastin time (aPTT) risk scores (39–42).

Yet challenges remain with developing PGS. Analyses suggest that only including SNPs reaching genome-wide significance in a PGS fails to capture much of the heritable variation and reduces the PGS’s prediction accuracy. However, deciding on a p-value threshold for including SNPs *a priori* is challenging. Recently, software has been developed which addresses this challenge, using summary statistics from GWAS to calculate a number of PGSs across a wide range of p-value thresholds for SNP inclusion and model fit statistics to determine the optimal threshold for predicting traits in a testing dataset, which is often less stringent than the genome-wide level (43).

To date, only a handful of studies have calculated PGSs for vitamin D concentrations, generally using only SNPs in genes that reached the stringent p-value threshold in existing vitamin D GWASs, therefore missing much of the genetic contribution to the phenotype (32,44–47). Given that several studies have reported genetic dependent response to vitamin D supplementation, PGSs hold predictive and preventive promise in relation to vitamin D concentrations (30,48,49).

The goal of the current study was to calculate ancestry-specific PGSs for 25(OH)D in individuals of European or African ancestry based on the results from a recent multi-ethnic GWAS meta-analysis (37), and to validate the PGS performance in an independent sample. Additionally, the proportion of SNP heritability captured by the PGS was quantified, using GCTA, and compared to that captured by the genome-wide significant SNPs.

## Materials and Methods

### GWAS summary statistics

The <u>TRANS</u>-ethni<u>C</u> <u>E</u>valuation of vitami<u>N</u> <u>D</u> GWAS consortium (TRANSCEN-D), performed the largest multi-ethnic vitamin D GWAS meta-analysis to date and included 13 cohorts (9 of African ancestry, 3 of Hispanic ancestry and the SUNLIGHT discovery cohort, a consortium of 15 European cohorts) (37). Here, ancestry-specific summary statistics from the African- and European-ancestry cohorts of TRANSCEN-D were leveraged for weighting of each SNP included in the PGS (37). This is referred to as the base dataset.

### Target/tuning dataset for calculation of PGS

Using weights from the base dataset, PGSs were developed in an ancestry-specific manner for subsets of European- and African-ancestry samples from the Atherosclerosis in Communities (ARIC) study, which contains European- and African-ancestry participants (50). These ARIC subsets are referred to as the target/tuning datasets. ARIC data were obtained through dbGaP Study Accession: phs000090.v4.p1. ARIC data were selected as the target/tuning dataset as they included both sexes and had dense genotyping, essential for development of a comprehensive and generalizable PGS. ARIC is a prospective epidemiologic study conducted across four United States sites: Wake Forest Baptist Medical Center, Winston-Salem, NC; University of Mississippi Medical Center, Jackson, MS; University of Minnesota, Minneapolis, MN; Johns Hopkins University, Baltimore, MD. ARIC includes 15,792 participants aged 45-64 at baseline, of which 9,086 have data required for this analysis (genomic data, 25(OH)D, age, sex, body mass index (BMI), location and month of blood draw) which were ascertained at ARIC visit 2 (1990-1992). Of these 9,086 participants, 7,178 are of European ancestry. A random sample of 1,000 participants were chosen from the 7,178 eligible European-ancestry participants for tuning the optimal PGS model, the remaining samples were used to validate the PGS. From the 1,908 eligible participants of African ancestry from ARIC, only data from 57 participants had not been included in the TRANSCEN-D meta-analysis; these were used as the base dataset. Therefore, to ensure independence between the base and target/tuning datasets only these 57 African-ancestry participants were selected into the target/tuning dataset.

### PGS validation cohort

PGS validation was done using data combined across participants from three multi-ethnic cohorts: ARIC, the Multi-ethnic Study of Atherosclerosis (MESA) and the Women’s Health Initiative (WHI), analyzed in an ancestry-specific manner. As above, ARIC provided data on 6,178 participants of European ancestry for the validation cohort. MESA is a prospective study of men and women ages 45-84 who were recruited by Columbia University, New York, NY; Johns Hopkins University, Baltimore, MD; Northwestern University, Chicago, IL; University of Minnesota, Minneapolis, MN; University of California at Los Angeles, Los Angeles, CA and Wake Forest University, Winston-Salem, NC. Serum 25(OH)D was measured at MESA exam 1 (July 2000-August 2002). MESA data were obtained through dbGaP Study Accession: phs000209.v13.p3. MESA provided data on 1,936 European- and 342 African-ancestry participants who maintained independence from TRANSCEN-D for the validation cohort. Women participating in WHI were recruited from 40 clinical centers in the United States. Serum 25(OH)D was measured as part of the Calcium and Vitamin D (CaD) Trial (51). WHI data were obtained through dbGaP Study Accession: phs000200.v11.p3. Participants were included if they had the minimum set of variables: genome-wide data, serum 25(OH)D, age, sex, BMI, and month of blood draw. WHI provided data on 455 European- and 700 African-ancestry participants for the validation cohort. Thus, together, in the validation cohort, the European-ancestry sample included 8,569 participants and the African-ancestry sample included 1,042 participants.

### Datasets used for heritability estimation

Heritability estimates were calculated using eligible participants (N=8,838) of both European (n=7,119) and African (n=1,719) ancestry from ARIC.

Participant consent was previously obtained for each study providing data; additionally, IRB approval was granted for this specific mega-analysis.

### Data Quality Control

Data cleaning for phenotypic data included winsorizing 25(OH)D in the MESA and WHI samples to account for outliers (52). In the WHI sample, participants with 25(OH)D values far above the maximum level of detection (150 ng/mL), none of which had extreme vitamin D intake (including supplement use) or sun exposure, were removed from the sample; this included 68 participants of European ancestry and 119 participants of African ancestry. All 25(OH)D values were log transformed to improve the normality of the distribution in each cohort.

Genotyping methods are available in Supplemental Table 1 and described in more detail elsewhere (53–57). In summary, QC removed: sex mismatches, samples and SNPs with high missingness (>5%), SNPs with low minor allele frequency (MAF<0.2%), and SNPs out of Hardy-Weinberg equilibrium (HWE<0.05/number of SNPs; Bonferroni adjusted cut-off). Datasets were imputed using the Michigan Imputation Server (58,59). European samples were imputed to the Haplotype Reference Consortium (HRC) and African samples were imputed to the Consortium on Asthma among African-ancestry Populations in the Americas (CAAPA) (59,60). Post imputation QC included: removing SNPs with a low quality score (<0.8) or MAF (<0.1%). Additionally, sample and SNP level missingness as well as HWE cutoffs were rechecked. Supplemental Figures 1-2 and Supplemental Table 1 give specifics on quality control for each cohort. QC was performed using PLINK v1.9 and vcfTools (61,62). Ancestry was determined by self-reported data and confirmed with principal components analysis (PCA) in PLINK using 1000 Genomes data as anchoring populations (63).

### Measurement of 25(OH)D

Blood 25(OH)D concentration was measured by the studies using different assay types. WHI used the DiaSorin LIASON chemiluminescence, while both MESA and ARIC used liquid chromatography-mass spectrometry (LCMS), which is considered the gold standard for 25(OH)D measurement (64,65). Vitamin D concentration [25(OH)D] was log transformed to improve normality of the distribution. To control for differences in vitamin D concentrations due to different assays, 25(OH)D concentrations were converted to z-scores within studies for combined cohort analyses.

### Measurement of vitamin D intake

Dietary data were collected via questionnaire. Each study used their own questionnaire. WHI used the Food Frequency Questionnaire supplemented with interview questions. ARIC and MESA both used their own implementation of a food intake questionnaire. From the questionnaire data, each study created a derived variable of typical vitamin D intake (measured in IU or mcg). All values were converted to IU for analysis. Additionally, WHI collected data on vitamin D supplement use at the same visit that 25(OH)D was assessed. The sum of vitamin D intake from food and supplements was calculated and used for supplemental and sensitivity analyses, otherwise dietary intake alone was used.

### Calculation of available UV radiation

Available UV radiation was calculated based on the month of blood draw and location; participants were assigned continuous available UV radiation values. Available UV radiation values assigned were an average UV-index for the month prior to blood draw (the relevant exposure period). UV data come from the National Weather Service Climate Prediction Center historical database. When available, UV radiation values corresponded to the exact location and year of the participant’s blood draw. When exact cities or years were not available, averages across nearby locations or years were used. See Supplemental Tables 2-4 for specific month, year and location values used. Descriptive statistics for available UV radiation values by site and month are also presented in Supplemental Tables 5-7; the UV radiation values ranged from 0.7 to 9.5 UV Index units.

### Determining optimal PGS

An optimal p-value cutoff and corresponding PGS were determined by calculating PGSs across a wide range of p-value thresholds and testing the association between the PGS and log[25(OH)D] in the target/tuning dataset. First, summary statistics were attained from TRANSCEN-D, the base dataset (1,37), which included SNPS with MAF > 0.01 and tested them for association with log[25(OH)D] using an additive genetic model adjusting for age, sex, BMI, UV index and principal components (PCs) 1-10. PGSs were then calculated in PRSice v2, which computes the sum of reference allele counts at each SNP weighted by the effect size (β) for that SNP from the TRANSCEN-D consortium (43). PGS weights came from ancestry-specific z-scores from TRANSCEN-D that were converted to betas with the deterministic relationship: β = *z*/(*sqrt*(2*p*(1 − *p*) * *N*)), where *p* is the allele frequency for the reference SNP (66). One tuning parameter in PGS development is the LD cutoff used for clumping to prevent SNPs in one correlated region from dominating the PGS. Here, PGSs were calculated in the target/tuning dataset using two different LD cut-offs, r^2^≥ 0.5 or ≥ 0.2, keeping the SNP with the strongest effect in the base dataset. SNPs in LD with one another were clumped, using the --clump-r2 option in PRSice v2. The LD cutoff that yielded the PGS that explains the most variance in 25(OH)D was used in downstream analyses. Given the small African-ancestry sample, a reference panel (remaining ARIC African-ancestry dataset n= 1,900) was used to determine LD.

To determine which set of SNPs to include in the PGS, SNPs at or below a given p-value threshold in the base dataset were included in the PGS and tested in a linear regression model for association with log[25(OH)D in the target/tuning dataset]. P-value thresholds from 5×10^−5^-0.5 were tested incrementing by 5×10^−5^ at each iteration (supplemental testing using a p-value threshold of 1.0 was performed). All testing was done using PRSice v2 (43). The threshold with the PGS explaining the most variance in log[25(OH)D] was selected as the most optimal PGS. R^2^, or the coefficient of determination, was calculated to measure the proportion of phenotypic variance explained by the model. Linear regression models were used to calculate the R^2^ of a given PGS while controlling for participant age, sex, BMI, available UV radiation, and PCs for ancestry. Five PCs were controlled for in the African-ancestry models and two PCs in the European-ancestry models, as determined based on the ‘elbow’ of cohort- and ancestry-specific scree plots. Sensitivity analysis was performed including dietary intake in the model, as dietary intake is a strong predictor of 25(OH)D concentrations. However, with the inclusion of dietary intake in the model, the optimal PGS (and p-value cutoff) remained the same for the European cohort, but reduced sample size substantially in the African-ancestry cohort. Therefore, to maintain sample size, dietary intake was not included in the model to determine the optimal p-value cutoff (Supplemental Table 8).

### PGS performance validation

PGS performance was validated in an ancestry-specific manner using participants in validation cohorts, which were combined cohorts of samples from ARIC, MESA, and WHI that maintained independence from TRANSCEN-D and PGS development samples. The PGS was applied to the participants in the validation dataset in accordance with the ancestry specific p-value cutoff. The relationship between PGS quantile and 25(OH)D was tested using a linear regression model controlling for age, sex, BMI, available UV radiation, and PCs for ancestry. Quantile plots were created depicting the relationship between PGS decile and 25(OH)D concentration. Sensitivity analyses were performed to ensure that the study design of the WHI CaD randomized control trial was not biasing the results.

### Supplemental PGS Analyses

The African-ancestry target/tuning cohort was small (n=57) due to limited genome-wide data in those of African ancestry. To explore if the small sample reduced prediction for those of African ancestry, a PGS was created from all independent SNPs (p-value cutoff = 1.0; r^2^ cutoff = 0.5) in the full independent sample of African-ancestry participants which maintained independence from TRANSCEN-D (n=1,099) (67). Additionally, to test the importance of ancestrally-matched base and target sets, this PGS was also created using European-ancestry GWAS summary statistics for weighting of the PGS.

### Heritability Estimation

Heritability estimates were calculated using GCTA v1.26 (68). Heritability was estimated several ways: (1) ancestry-specific overall SNP heritability, (2) ancestry-specific SNP heritability of the PGS (where sample size allowed) and (3) ancestry-specific SNP heritability of previous replicated GWAS findings in *CYP2R1, CYP24A1, DHCR7, and GC* (1,34,37). In each case, the model was adjusted for age, sex, BMI, available UV, and dietary vitamin D intake.

SNP heritability estimates were calculated using all genotyped and imputed SNPs for both the European and African ancestry populations from ARIC; this was 8,315,761 and 9,335,785 SNPS, respectively. Partitioned heritability estimates were discerned paralleling methodology described by the SUNLIGHT consortium (36). To estimate heritability captured by the PGS, heritability was calculated twice; once using the clumped set of SNPs used to determine the PGS (228,867 SNPs for European ancestry and 850,697 for African ancestry) and a second time using the clumped set of SNPs with SNPs included in the PGS removed (228,526 SNPs for European ancestry and 818,428 for African ancestry). The difference in heritability estimates between these two models was the heritability explained by the PGS. Heritability could not be directly calculated from the SNPs in the PGS because one of the assumptions made by the GCTA modeling is an average null effect of the SNPs on the outcome. Of note, the African-ancestry sample was too small for this analysis to be valid, so heritability attributed to the PGS was only calculated in those of European ancestry. In discerning the heritability captured by previous replicated GWAS studies, heritability was calculated using a reduced set of SNPs: the full genotyped and imputed set with top GWAS findings (and SNPs in the surrounding LD block) removed (36,69). The difference between this estimate and the overall heritability estimates was the heritability attributed to previous replicated GWAS findings. Additionally, a second heritability estimate was calculated that included novel findings. This included SNPs from *AMDHD1* and *SEC23A* in those of European ancestry and SNPs from *KIF4B, HTR2A* and *ANO6/ARID2* in those of African ancestry (36,37). Table 1 summarizes the SNPs and LD blocks removed in each scenario. LD block size was determined using the Plots mode of the SNAP tool by the Broad (69). All models were fit separately for European and African ancestry samples.

**Table 1:**
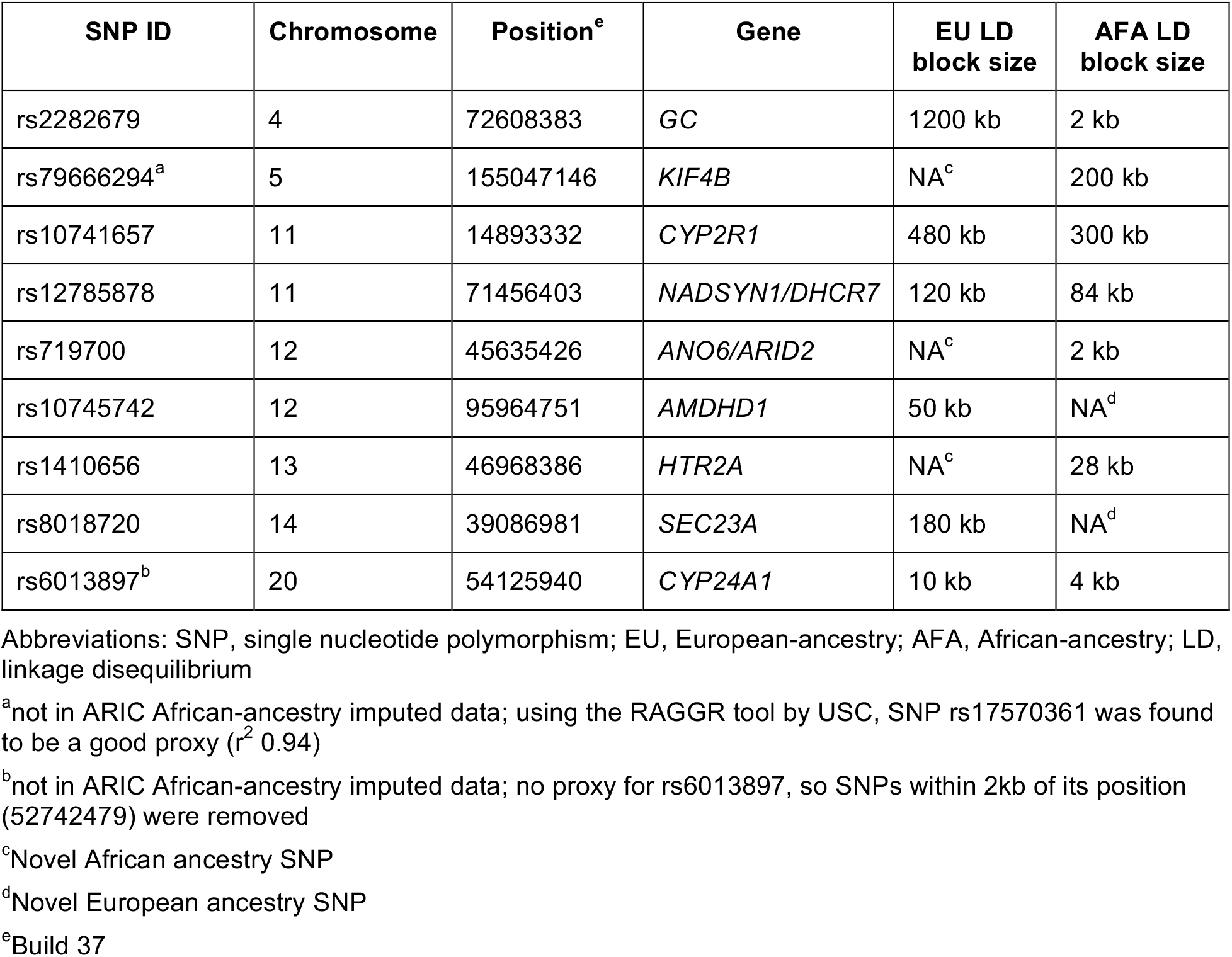
Previous GWAS SNPs

## Results

### Determining the optimal PGS

Table 2 shows sample characteristics for each analysis. Table 3 shows statistics for the best performing PGS for each ancestry in the target/tuning and validation datasets while controlling for age, sex, BMI, available UV radiation, and PCs for ancestry. In both ancestries, the PGS using the LD cut-off of 0.5 was more strongly associated with and explained more of the variance in log[25(OH)D] than did the PGS using the LD cut-off of 0.2 (Supplemental Table 9). Therefore, this was the LD cutoff utilized going forward. In the European-ancestry analyses, the optimal PGS explained 1.4% of the variance in log[25(OH)D] (p=9.3×10^−5^) in the target/tuning dataset and 1.0% of the variance (p=1.1×10^−23^) in the validation cohort. In the African-ancestry analyses, the PGS explained 2.9% of the variance in log[25(OH)D] (p=0.11) in the target/tuning dataset and 0.2% of the variance (p=0.15) in the validation cohort. Of note, the optimally performing PGS in the African-ancestry target/tuning dataset contained many more SNPs than that from the European-ancestry dataset, mostly due to the less stringent p-value cutoff, but also because a larger number of SNPs remained post clumping (850,697 vs 228,867) due to smaller LD blocks in the African-ancestry sample and more input SNPs from the TRANSCEN-D summary statistics (8.4 million in the African-ancestry vs 1.2 million in the European-ancestry sample). Figure 1 depicts the results visually, where a taller bar corresponds to a larger percent of the phenotypic variance explained by the PGS.

**Figure 1.**
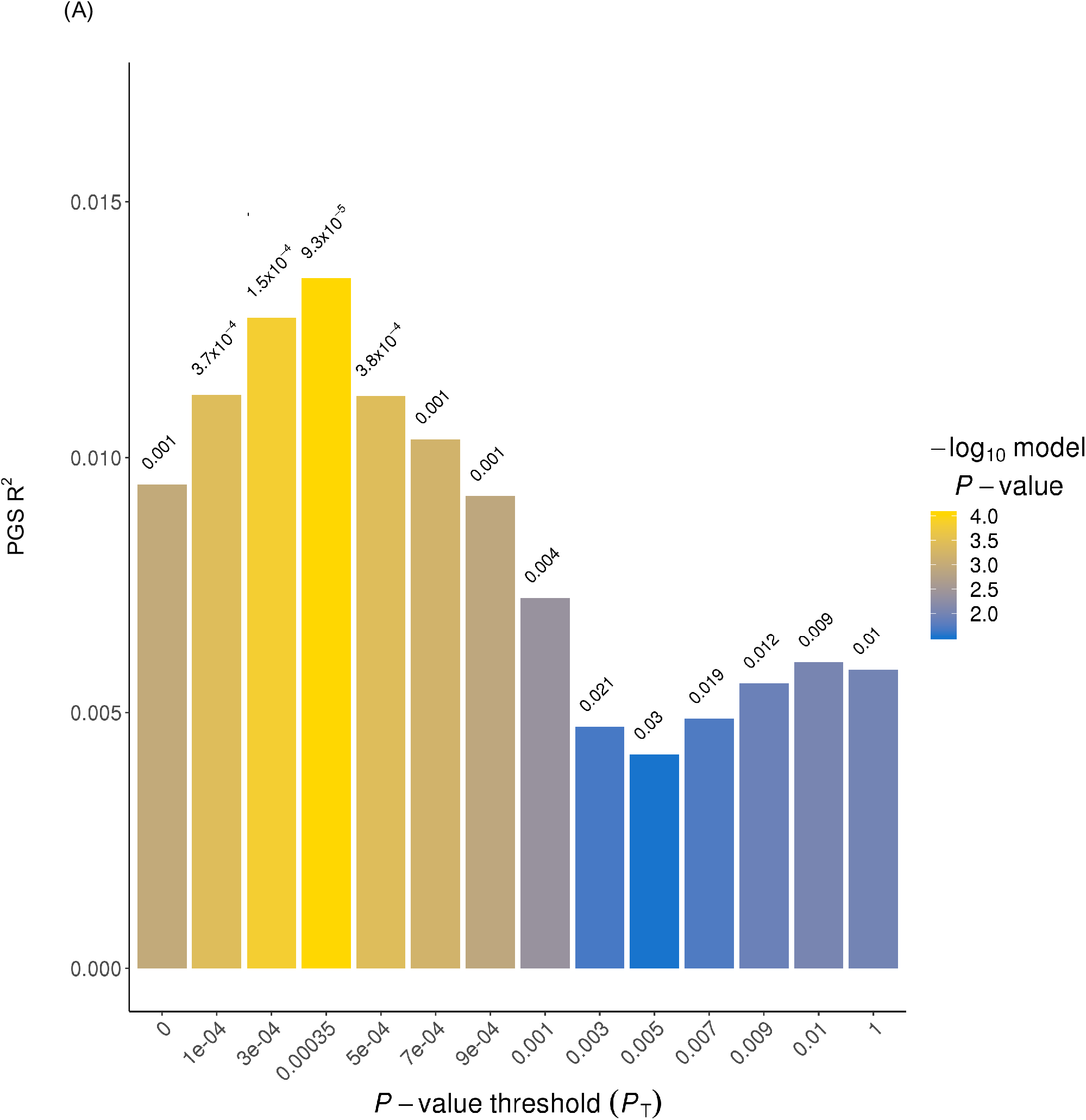
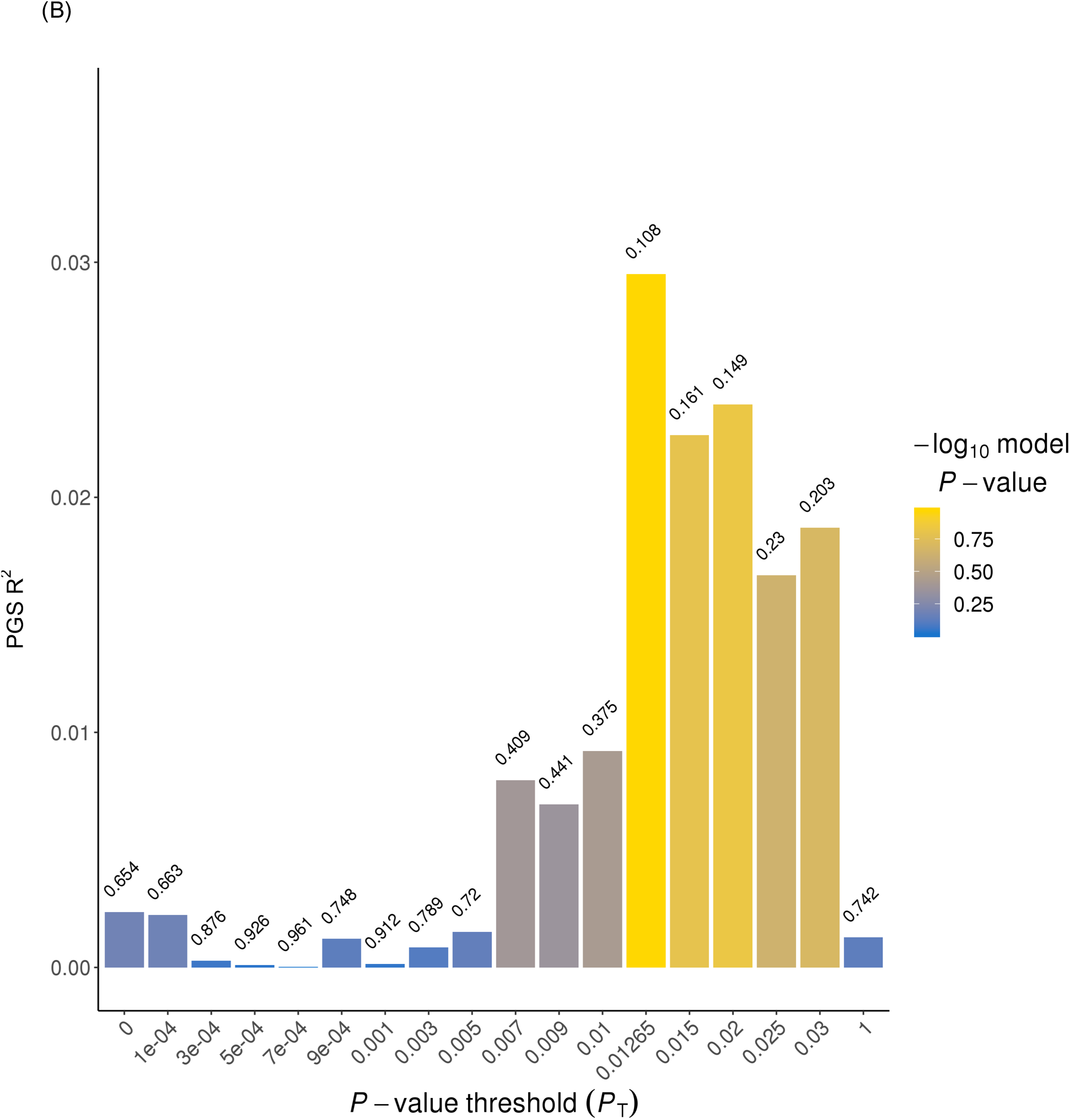
The polygenic score (PGS) performance in those of European or African ancestry. The x-axis displays selected p-value thresholds for single nucleotide polymorphisms (SNPs) included in the PGS. The y-axis displays the proportion of phenotypic variance captured by the PGS. Yellow bars correspond to a strong association (more significant p-value) between the PGS and log(25(OH)D than do blue bars. In panel A, the most optimally performing PGS has the tallest bar (p-value threshold = 0.00035) and captures 1.4% of the variance in log[25(OH)D]. In panel B, the most optimally performing PGS has the tallest bar (p-value threshold = 0.01265) and captures 2.9% of the variance in log[25(OH)D].

**Table 2.** Sample characteristics

| Sample | Metric | European-ancestry set | African-ancestry set |
| --- | --- | --- | --- |
| **PGS development sample (from ARIC)** | **Sample Size** | 1,000 | 57 |
|  | **% Female** | 53.2% | 49.1% |
|  | **Mean Age (SD) [years]** | 57.1 (5.7) | 55.6 (6.2) |
|  | **Mean BMI (SD) [kg/m<sup>2</sup>]** | 27.3 (4.8) | 28.6 (5.7) |
|  | **Mean Available UV radiation (SD) [units]** | 5.0 (2.5) | 7.1 (2.4) |
|  | **Mean Vitamin D Intake (SD) [IU]** | 219.2 (135.2) | 221.2 (137.3) |
|  | **Mean 25(OH)D (SD) [ng/ml]** | 25.7 (8.7) | 20.9 (7.8) |
| **PGS validation sample (from ARIC, MESA and WHI)<sup>a</sup>** | **Sample Size** | 8,569 | 1,042 |
|  | **% Female** | 56.1% | 84%<sup>b</sup> |
|  | **Mean Age (SD) [years]** | 58.9 (7.7) | 62.0 (8.5) |
|  | **Mean BMI (SD) [kg/m<sup>2</sup>]** | 27.5 (5.0) | 30.8 (6.3) |
|  | **Mean Available UV radiation (SD) [units]** | 4.9 (2.5) | 5.4 (2.5) |
|  | **Mean Vitamin D Intake (SD) [IU]** | 172.3 (157.0) | 99.7 (126.1) |
|  | **Mean 25(OH)D (SD) [ng/ml]** | 26.5 (9.7) | 19.2 (13.6) |
| **Heritability estimation sample (from ARIC)** | **Sample Size** | 7,119 | 1,719 |
|  | **% Female** | 53.6% | 63.5% |
|  | **Mean Age (SD) [years]** | 57.1 (5.7) | 56.4 (5.8) |
|  | **Mean BMI (SD) [kg/m<sup>2</sup>]** | 27.3 (4.8) | 30.2 (6.2) |
|  | **Mean Available UV radiation (SD) [units]** | 5.0 (2.5) | 6.9 (2.3) |
|  | **Mean Vitamin D Intake (SD) [IU]** | 222.8 (144.4) | 215.7 (150.3) |
|  | **Mean 25(OH)D (SD) [ng/ml]** | 25.9 (8.8) | 19.1 (7.1) |

Abbreviations: PGS, polygenic score; SD, standard deviation; BMI, body mass index; UV, ultraviolet
<sup>a</sup>MANOVA global test (performed in SAS (version 9.4) revealed differences in one or more variables by cohort, therefore cohort was adjusted for in all models that included multiple cohorts
<sup>b</sup>Notably greater % Female than other sets, because this set includes independent samples in MESA (n=342) and WHI (n=700)

**Table 3:** Performance of optimal PGS in each ancestry

| Ancestry | p-value cut-off | # SNPs | Cohort | PGS $R^2$ (model $R^2$) | p-value<sup>a</sup> |
| --- | --- | --- | --- | --- | --- |
| European | 0.00035 | 341 | Tuning (n=1,000) | 0.014 (0.14) | $9.3 \times 10^{-5}$ |
| | | | Validation (n=8,569) | 0.0098 (0.17) | $1.1 \times 10^{-23}$ |
| African | 0.01265 | 32,269 | Tuning (n=57) | 0.029 (0.49) | 0.11 |
|  |  |  | Validation (n=1,042) | 0.002 (0.03) | 0.15 |
| African | 1.0 | NA<sup>b</sup> | Full-Independent (n=1,099) | 0.003 (0.04) | 0.05 |

Abbreviations: SNPs, single nucleotide polymorphism; PGS, polygenic score
<sup>a</sup>p-value for association between PGS and log[25(OH)D]
<sup>b</sup>this PGS was created via mega-analysis of three cohorts; # SNPs varies by cohort

In supplementary analyses for the African-ancestry cohort, the PGS using the full African-ancestry sample that was independent from TRANSCEN-D (n=1,099; p-value cutoff = 1.0; r^2^ cutoff = 0.5) explained more variance (0.31% vs 0.2%) and had a stronger association (p=0.0545 vs 0.15) than the optimal PGS determined from PRSice using the small tuning cohort (n=57) and larger validation cohort (n=1042), therefore this PGS was chosen as most optimal and used moving forward. In additional analyses to test the importance of ancestrally-matched base and target sets, the PGS (p-value cutoff = 1.0; r^2^ cutoff = 0.5) developed from European-ancestry TRANSCEN-D summary statistics only explained 0.14% of the variance (p=0.1897; worse performance than the PGS developed from African-ancestry TRANSCEN-D summary statistics that explained 0.31 % of the variance in 25(OH)D [p=0.0545]). Results are shown in Table 3 and Supplemental Table 10.

After the optimal, ancestry-specific PGS was discerned, the relationship between the PGS and 25(OH)D was investigated using ancestry-specific combined cohorts of samples from ARIC, MESA and WHI, which maintained independence from the TRANSCEN-D (and tuning, for those of European ancestry) samples. Characteristics of these samples are summarized in Table 2. Figures 2 and 3 show ancestry-specific plots for 25(OH)D by decile of the PGS. In general, those with greater genetic risk (lower PGS and quantile) have lower 25(OH)D concentrations. For a clinically-based interpretation, in the European validation cohort (Figure 2, n=8,569), those with the lowest PGS have vitamin D concentrations 3.0 ng/ml lower than those with the highest PGS (p=3.2×10^−13^). Figure 3 shows the trend for those of African ancestry (n=1,099; combined tuning and validation samples from Table 2); those with the lowest PGS have vitamin D concentrations 2.8 ng/ml lower than those with the highest PGS (p=0.0463). Results from the PGS determined using the separate tuning and validation cohorts are included in Supplemental Figure 3.

**Figure 2.**
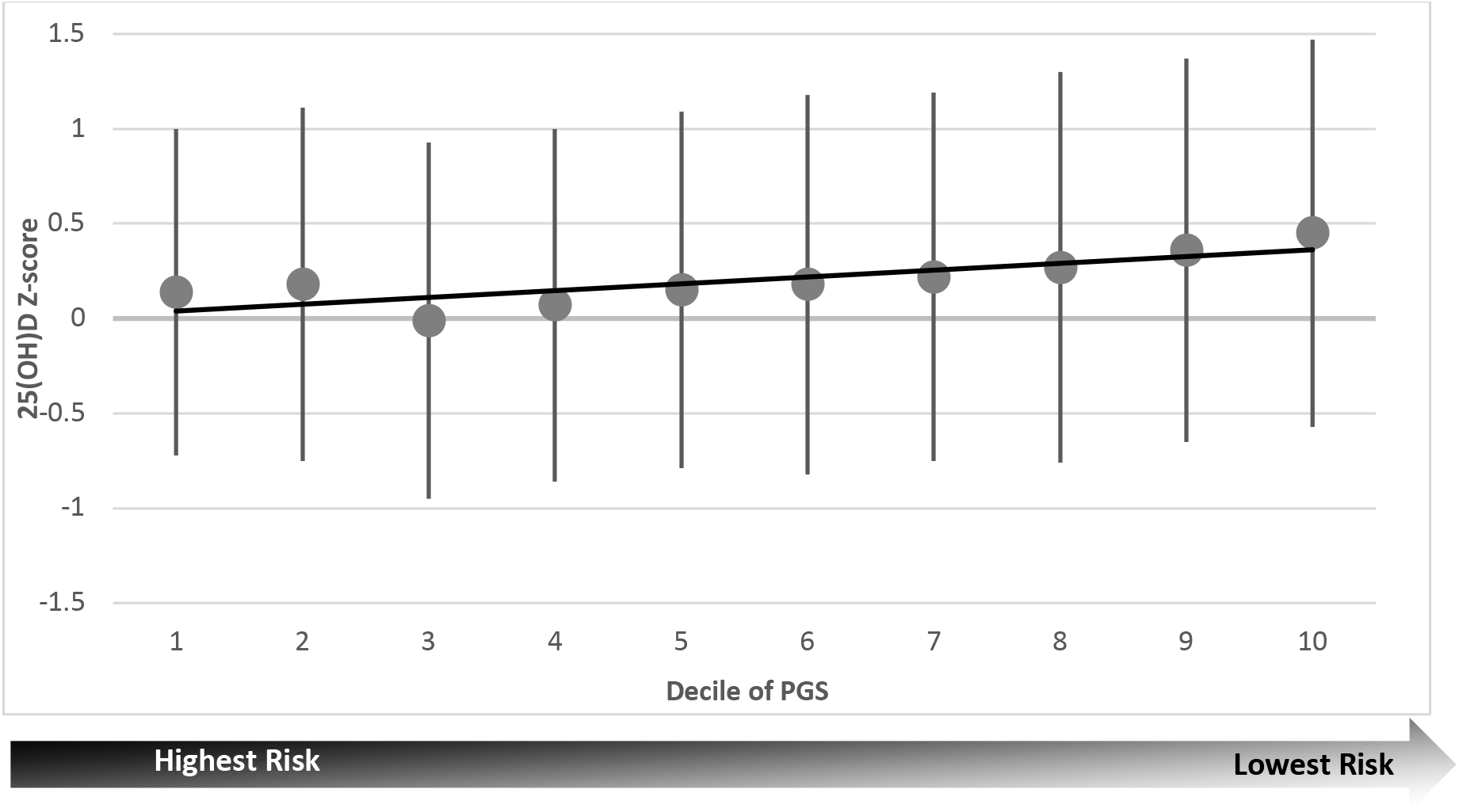
Visual representation of the association between polygenic score (PGS) decile and normalized vitamin D concentrations in those of European ancestry. The x-axis is the PGS decile, where lower decile means more risk of low vitamin D concentrations The y-axis is vitamin D concentrations (normalized for comparison between cohorts). Panel A is a plot for the subset of ARIC samples used to discern the optimal p-value threshold for the PGS (n=1000); panel B is a plot for the remaining independent samples (n=8,569). While the exact trend varies by plot, the general trend is that when the PGS decreases (i.e. higher genetic risk) 25(OH)D concentrations decrease. In panel A, moving from the lowest risk to the highest risk decile decreases vitamin D concentrations by 4.0 ng/ml. In panel B, moving from the lowest risk to the highest risk decile decreases vitamin D concentrations by 3.0 ng/ml.

**Figure 3.**
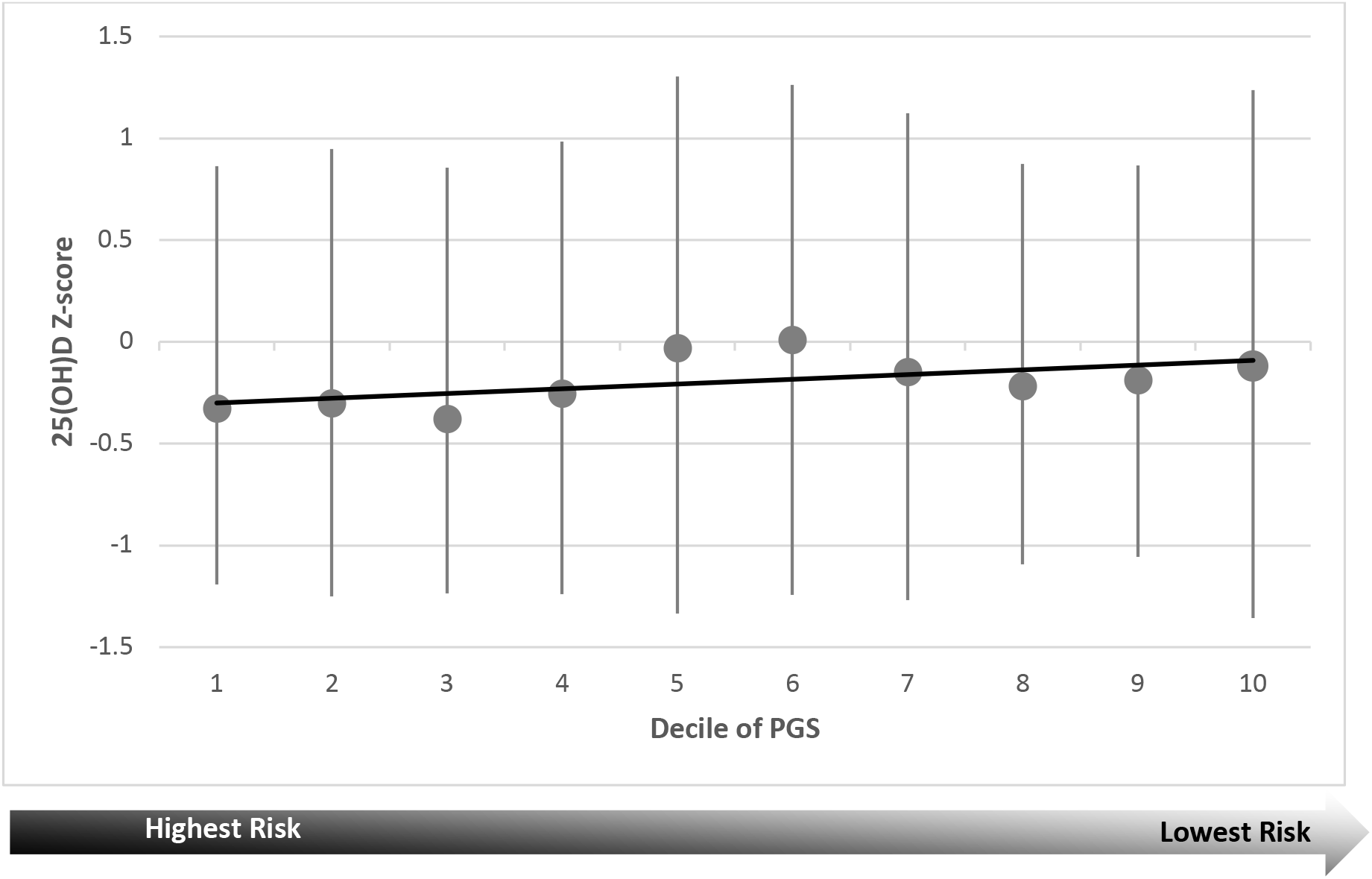
Visual representation of the association between polygenic score (PGS) decile and normalized vitamin D concentrations in those of African ancestry (n=1,099). The x-axis is the PGS decile, where lower decile means more risk of low vitamin D concentration. The y-axis is vitamin D concentrations (normalized for comparison between cohorts). The trend is that when the PGS decreases (i.e. higher genetic risk) 25(OH)D concentrations decrease. Moving from the lowest risk to the highest risk quintile decreases vitamin D concentrations by 2.8 ng/ml (p=0.0463).

Sensitivity analyses showed there was no significant difference in 25(OH)D concentration between participants on the treatment arm compared to the placebo arm in the participants from WHI. Additionally, there was no significant difference in PGS-25(OH)D trend in WHI compared to the other cohorts.

### Heritability estimation

Participant characteristics are summarized in Table 2. Overall and stratified SNP heritability estimates for those of European or African ancestry are summarized in Figure 4 and Supplemental Figure 4, respectively. SNP heritability is higher in the African-ancestry cohort compared to the European-ancestry cohort (32% vs 22%; standard errors 17.8 and 5.2, respectively (p=0.49). In those of European ancestry, the PGS accounts for 17.1% (3.7/21.6) of the SNP heritability of 25(OH)D concentrations and previous replicated GWAS findings (i.e., SNPs from *CYP2R1, CYP24A1, DHCR7* and *GC*) account for 6.9% (1.5/21.6) of the total SNP heritability (1,34,37). In those of African ancestry, these same top GWAS findings accounted for only 1.6% (0.5/32.2) of the total SNP heritability. Heritability accounted for by previous GWAS findings remained unchanged when ancestry-specific novel findings were included in the heritability estimations (1,34,36,37). African-ancestry sample size was too small to calculate heritability accounted for by the PGS.

**Figure 4.**
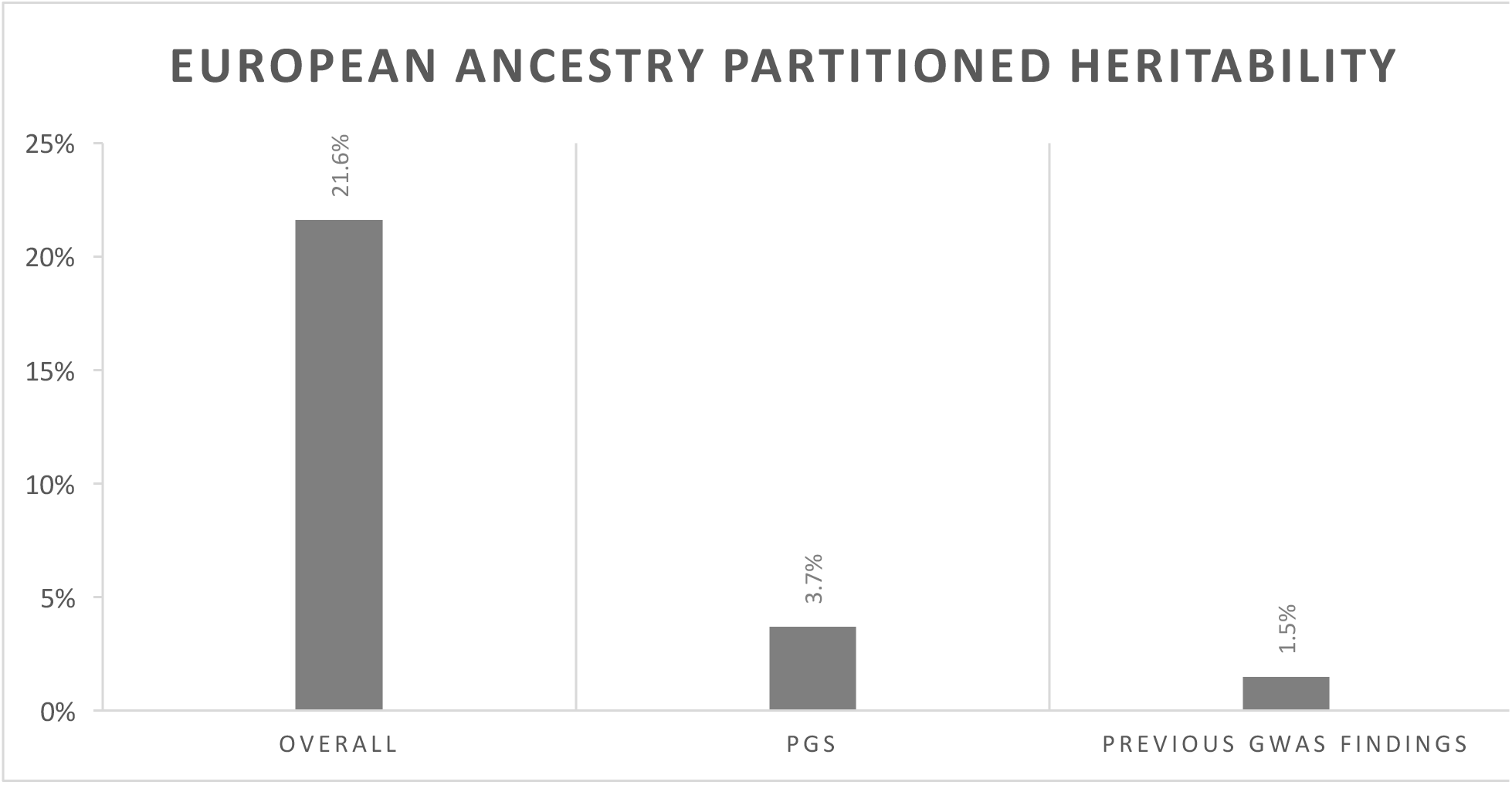
Overall single nucleotide polymorphism (SNP) heritability, polygenic score (PGS) SNP heritability and replicated genome-wide assocation study (GWAS) SNP heritability in those of European ancestry. Where sample allowed for calcuation, the PGS explains more heritability than previous replicated GWAS findings, albeit leaving much of the heritability unexplained.

## Discussion

Vitamin D inadequacy is a pervasive health problem, with a strong genetic basis. However, to date, much of the heritability of 25(OH)D remains unexplained. Furthermore, there is a tremendous gap in the research carried out in minority ancestries compared to European ancestry. Filling these knowledge gaps is critical in preventive care to manage 25(OH)D concentration, especially as we move towards precision medicine, and development of an ancestry-specific PGS is one way to address these gaps. To date, across all phenotypes, most PGS have been calculated in those of European ancestry. A handful of studies have begun to explore ancestry-specific PGS, however, none of these approaches utilize an entirely ancestry-specific approach as was undertaken here (70–72). Given the underlying genetic difference between ancestries (i.e., different LD patterns and allele frequencies), an ancestry-specific approach is more appropriate. Calculating PGSs using GWAS summary statistics from an ancestry-matched population accounts for differences in linkage disequilibrium (LD) and allele frequencies that exist between ancestral groups leading to differences in allele effect sizes, which are used as weights in the PGS calculation. Here, optimal PGSs were discerned and validated in an ancestry-specific manner. Heritability explained by the PGS and previous GWAS findings was compared to overall SNP heritability.

The relationship between the PGS and 25(OH)D concentrations was consistent across ancestries in the validation cohorts, albeit modest variance was explained by the PGS; those with the lowest PGS (most risk) had the lowest 25(OH)D concentrations. Moving from the highest to lowest quantile changed 25(OH)D concentrations by 2.8-3.0 ng/ml, a statistically significant and clinically meaningful difference. One study reported that for each additional 100 IU of vitamin D consumed, serum 25(OH)D levels increased by 0.6 ng/ml (73). Using this conversion, compared to those with lowest genetic risk, those with highest genetic risk could require an additional 467 to 500 IU of vitamin D to maintain comparable levels. While the small sample used to determine the p-value cut-off in those of African ancestry could have led to overfitting of the model, the consistent direction of effect between PGS quantile and 25(OH)D concentrations suggests clinical utility for a 25(OH)D PGS to inform vitamin D supplementation in those with high genetic risk for 25(OH)D inadequacy.

The portion of phenotypic variance explained by the PGS was modest due to many concurrent influences. First, the PGS did not include rare variants (MAF < 0.01) as they were removed from the base set (TRANSCEN-D). Common SNPs account for only a small proportion of genetic variance in complex traits (38). Future PGSs that include rare variants will likely account for a greater portion of the variance. Additionally, the variance that the PGS can capture is limited by the input SNPs. In the best-case scenarios (i.e. densest chips), the overlap between the SNPs in the base and target datasets was 3,520,049 and 1,026,643 SNPs, for African and European ancestries, respectively. While over 1 million SNPs can be very informative, much of the genome was not included. Thirdly, PRSice implements clumping which keeps only the SNP with the strongest association for SNPs in LD (r^2^ >0.5 used here) in any given 500kb window, thus reducing the maximum variability that could be captured by a PGS.

Supplementary analyses performed in the full independent sample of African ancestry participants (n=1,099) demonstrated (1) the importance of a large tuning sample and (2) the importance of ancestrally-matched base and target sets. Comparing the results from the small tuning sample (n=57) and the analysis using the full independent sample of African-ancestry participants (n=1,099), more variance, 0.3% compared to 0.2%, was accounted for, and the association between PGS and 25(OH)D was stronger (p=0.0545 vs 0.15) using the PGS developed in the full sample (n=1,099). Additionally, variance accounted for dropped to 0.14% (compared to 0.31%) and the PGS-25(OH)D association became non-significant (p=0.19) when using mismatched summary statistics, reiterating the importance of ancestry-specific analyses.

Not surprisingly, the heritability investigation provided further evidence that PGSs using a less stringent p-value threshold account for a higher portion of the heritability than genome-wide significant SNPs from previous GWAS. Here, the PGS explained more of the SNP heritability than did previous GWAS findings, 17.1% compared to 6.9% in those of European ancestry (sample size was too small for PGS heritability calculations in those of African ancestry). However, neither the PGS nor previous GWAS findings explain a large portion of total SNP heritability, promoting the need for genetic studies with larger sample sizes and more dense SNP data that include low frequency variants to fully understand the genetic determinants of 25(OH)D concentrations and, therefore, inform the most effective vitamin D supplementation practices.

This study calculated a PGS with a moderate R^2^, a consistent relationship to 25(OH)D concentrations and that explained more heritability of 25(OH)D than previous GWAS findings, reiterating the importance of capturing genetic risk by PGS which can be used for clinical predictions. Additionally, this study contributes in-depth multi-ethnic investigation into 25(OH)D heritability by ancestry, teasing apart genetic underpinnings of 25(OH)D concentrations. However, the study does come with some limitations. To maintain independence from TRANSCEN-D, which provided ancestry-specific weights for the PGSs, the sample size used in this analysis was relatively small, especially for the African-ancestry cohort. The sample size issues experienced for the African-ancestry cohort emphasize the importance of obtaining more diverse samples (i.e. in initiatives like All of Us) (74). Through the TRANSCEN-D GWAS meta-analysis and the analysis here, nearly all of the publicly available African-ancestry samples with relevant data have been exhausted and sample sizes for other racial/ethnic groups remain limited. The limited amount of diverse data led to a small African-ancestry training set. A small training set limits discrimination of risk groups which could explain the less significant findings for those of African ancestry (42). Furthermore, the genotyping performed did not capture rare variants, limiting the variance that could be captured by the PGS. While GCTA allows for the calculation of heritability in non-related participants, which avoids overestimation due to shared environment, it only accounts for additive SNP effects, potentially underestimating total heritability which also could include gene-by-gene interactions. Finally, while adjusting for available UV radiation is more precise than season, it is not a perfect proxy for time spent outside and does not consider the amount of skin exposed, sunscreen use or skin pigmentation. These limitations leave room for future studies and replication that should be performed. For example, future PGSs could be developed implementing the recent cross-prediction method developed by Mak et,al; this method allows and corrects for overlap between the base and target dataset that would have allowed for a much larger African-ancestry sample (75). Additionally, in the future, the PGS could be utilized as an independent variable to predict health outcomes.

## Conclusion

This study showed that PGSs are a powerful predictive tool for determining 25(OH)D concentrations. Given the association between the optimal PGS and 25(OH)D concentrations, PGSs could be leveraged for personalized vitamin D supplementation, which could prevent the negative downstream effects of 25(OH)D inadequacy. Additionally, through an in-depth investigation of 25(OH)D SNP heritability, it was shown that the PGS explains more heritability than do GWAS findings to date. This provides additional evidence that many SNPs that function through small effect sizes influence 25(OH)D concentrations, yielding further understanding of the genetic architecture of 25(OH)D. However much of the heritability remains to be explained, therefore, more research is warranted along the quest to effectively and efficiently preventing 25(OH)D inadequacy through personalized supplementation.

**Appendix**

## Acknowledgments

We thank the researchers and participants of Atherosclerosis in Communities (ARIC), Multi-Ethnic Study of Atherosclerosis and the Women’s Health Initiative (WHI) for making this data publicly available.

## ARIC

The Atherosclerosis Risk in Communities Study is carried out as a collaborative study supported by National Heart, Lung, and Blood Institute contracts (HHSN268201100005C, HHSN268201100006C, HHSN268201100007C, HHSN268201100008C, HHSN268201100009C, HHSN268201100010C, HHSN268201100011C, and HHSN268201100012C). The authors thank the staff and participants of the ARIC study for their important contributions.

Funding for GENEVA was provided by National Human Genome Research Institute grant U01HG004402 (E. Boerwinkle). Vitamin D assays in ARIC were funded by R01 HL103706 (P. Lutsey).

## MESA

MESA and the MESA SHARe project are conducted and supported by the National Heart, Lung, and Blood Institute (NHLBI) in collaboration with MESA investigators. Support for MESA is provided by contracts HHSN268201500003I, N01-HC-95159, N01-HC-95160, N01-HC-95161, N01-HC-95162, N01-HC-95163, N01-HC-95164, N01-HC-95165, N01-HC-95166, N01-HC-95167, N01-HC-95168, N01-HC-95169, UL1-TR-000040, UL1-TR-001079, UL1-TR-001420, UL1-TR-001881, and DK063491.

The MESA CARe data used for the analyses described in this manuscript were obtained through Genetics (accession numbers). Funding for CARe genotyping was provided by NHLBI Contract N01-HC-65226.

Funding support for the Vitamin D dataset was provided by grant HL096875

## WHI

The WHI program is funded by the National Heart, Lung, and Blood Institute, National Institutes of Health, U.S. Department of Health and Human Services through contracts HHSN268201600018C, HHSN268201600001C, HHSN268201600002C, HHSN268201600003C, and HHSN268201600004C. This manuscript was not prepared in collaboration with investigators of the WHI, has not been reviewed and/or approved by the Women’s Health Initiative (WHI), and does not necessarily reflect the opinions of the WHI investigators or the NHLBI.

WHI PAGE is funded through the NHGRI Population Architecture Using Genomics and Epidemiology (PAGE) network (Grant Number U01 HG004790). Assistance with phenotype harmonization, SNP selection, data cleaning, meta-analyses, data management and dissemination, and general study coordination, was provided by the PAGE Coordinating Center (U01HG004801-01).

Funding support for WHI GARNET was provided through the NHGRI Genomics and Randomized Trials Network (GARNET) (Grant Number U01 HG005152). Assistance with phenotype harmonization and genotype cleaning, as well as with general study coordination, was provided by the GARNET Coordinating Center (U01 HG005157). Assistance with data cleaning was provided by the National Center for Biotechnology Information. Funding support for genotyping, which was performed at the Broad Institute of MIT and Harvard, was provided by the NIH Genes, Environment and Health Initiative [GEI] (U01 HG004424).

The datasets used for the analyses described in this manuscript were obtained from dbGaP at http://www.ncbi.nlm.nih.gov/sites/entrez?db=gap through dbGaP accession phs000200.v11.p3.

Funding for WHI SHARe genotyping was provided by NHLBI Contract N02-HL-64278.

KEH was supported by an NLM training grant to the Computation and Informatics in Biology and Medicine Training Program (NLM 5T15LM007359). Computational resources were supported by a core grant to the Center for Demography and Ecology at the University of Wisconsin-Madison (P2C HD047873).

