## Supplemental Materials for "Ancestry-specific polygenic scores and SNP heritability of 25(OH)D in African- and European-ancestry populations"

Hatchell, et al.

### Supplemental Tables and Figures

Supplemental Table 1: Genotyping information and quality control by cohort

| Cohort | SNP Array | n | SNP quality control | | | | Sample quality control | |
|---|---|---|---|---|---|---|---|---|
| | | | Call rate | Minor Allele Frequency | Hardy-Weinberg Equilibrium (post-imputation) | # SNPs Passing QC | Call rate | Exclusions |
| ARIC | AffymetrixGenome-WideHuman SNPArray 6.0 | African ancestry: 1,908<br>European ancestry: 7,178 | 95% | 0.002 | African Ancestry: $6x10^{-8}$ ($5x10^{-9}$)<br>European Ancestry: $6x10^{-8}$ ($6x10^{-9}$) | African ancestry: 9,335,785<br>European ancestry: 8,315,761 | 95% | sex mismatch, relatedness, chromosomal abnormalities |
| MESA | Affymetrix 50K gene-focused molecular imprinted polymer array (CVDSNP55v1_A) | African ancestry: 1,176<br>European ancestry: 1,936 | 95% | 0.002 | African Ancestry: $1x10^{-6}$ ($2x10^{-7}$)<br>European Ancestry: $1x10^{-6}$ ($1x10^{-7}$) | African ancestry: 309,712<br>European ancestry: 455,155 | 95% | sex mismatch, relatedness |
| WHI (Child6 NHLBI cohort) | AffymetrixGenome-WideHuman SNPArray 6.0 | African ancestry<br>Consent group 1: 65<br>Consent group 2: 572 | 95% | 0.002 | Consent group 1: $6x10^{-8}$ ($5x10^{-9}$)<br>Consent group 2: $6x10^{-8}$ ($5x10^{-9}$) | Consent group 1: 9,551,098<br>Consent group 2: 9,997,380 | 95% | sex mismatch, relatedness, race mismatch |
| WHI (Child7 GARNET cohort) | Illumina HumanOmni1-Quad v1-0 B | European ancestry:<br>Consent group 1: 86<br>Consent group 2: 443 | 95% | 0.002 | Consent group 1: $5x10^{-8}$ ($5x10^{-9}$)<br>Consent group 2: $5x10^{-8}$ ($4x10^{-9}$) | Consent group 1: 9,2037,621<br>Consent group 2: 9,722,526 | 95% | sex mismatch, relatedness, chromoso anomalies, race mismatch |
| WHI (Child 9 PAGE cohort) | Illumina MEGA Consortium 15063755 B2 array | African ancestry: 63 | 95% | 0.002 | $3x10^{-8}$ ($5x10^{-9}$) | African ancestry: 9,641,566 | 95% | sex mismatch, relatedness, chromoso anomalies, race mismatch |

Abbreviations: SNP, single nucleotide polymorphism; QC, quality control

[a]All African-ancestry samples were imputed to CAAPA using Michigan Imputation Server

[b]All European-ancestry samples were imputed to HRC r1.1 2016 using Michigan Imputation Server

[c]Sample sizes reported here reflect entire eligible cohort, actual numbers used differ to maintain independence where necessary

Supplemental Figure 1**:** Pre-imputation quality-control process

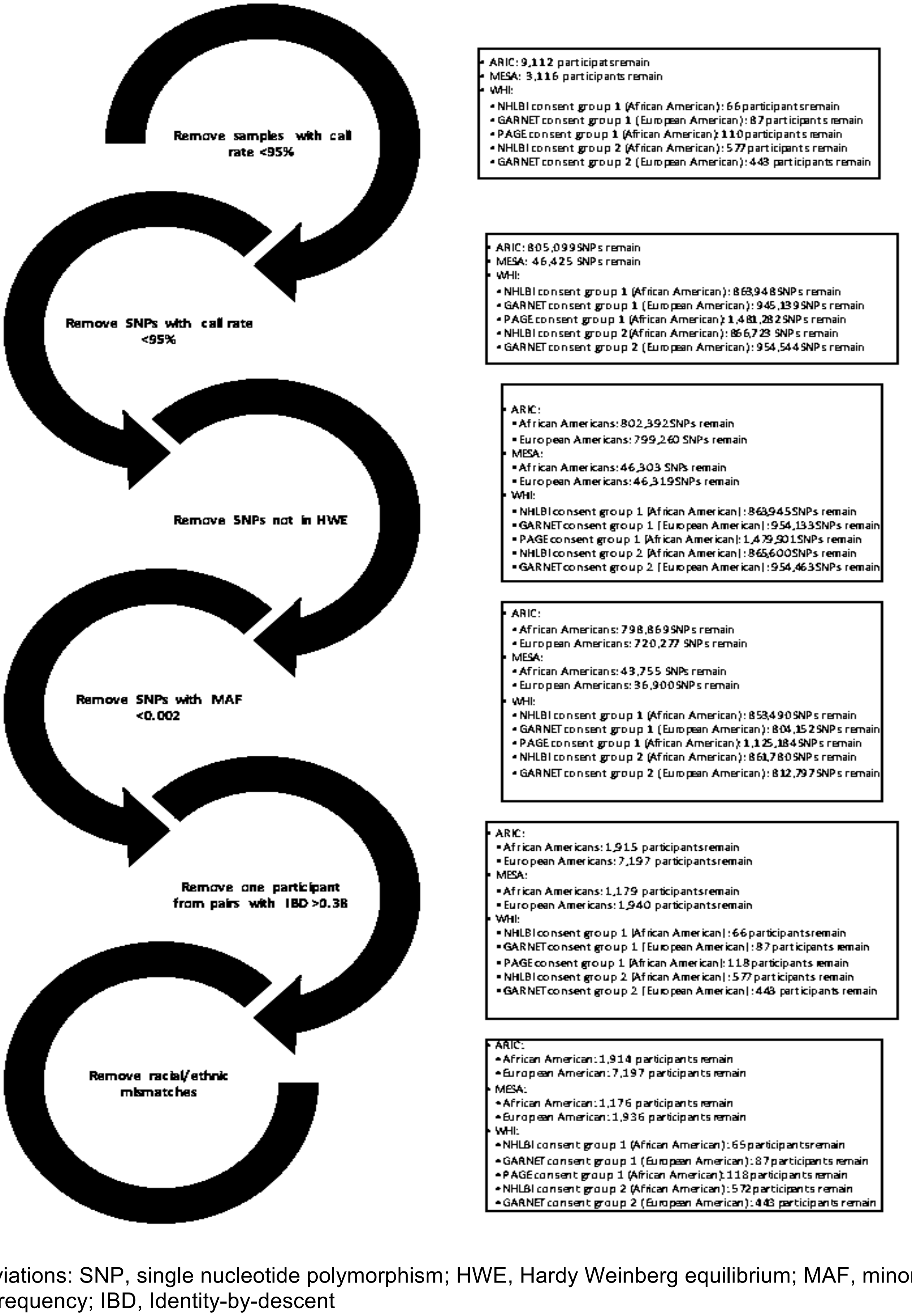


Abbreviations: SNP, single nucleotide polymorphism; HWE, Hardy Weinberg equilibrium; MAF, minor allele frequency; IBD, Identity-by-descent

Supplementary Figure 1 shows pre-imputation quality control steps and cutoffs used as well as corresponding sample sizes at each step.

Supplementary Figure 2: Quality-Control process starting at imputation

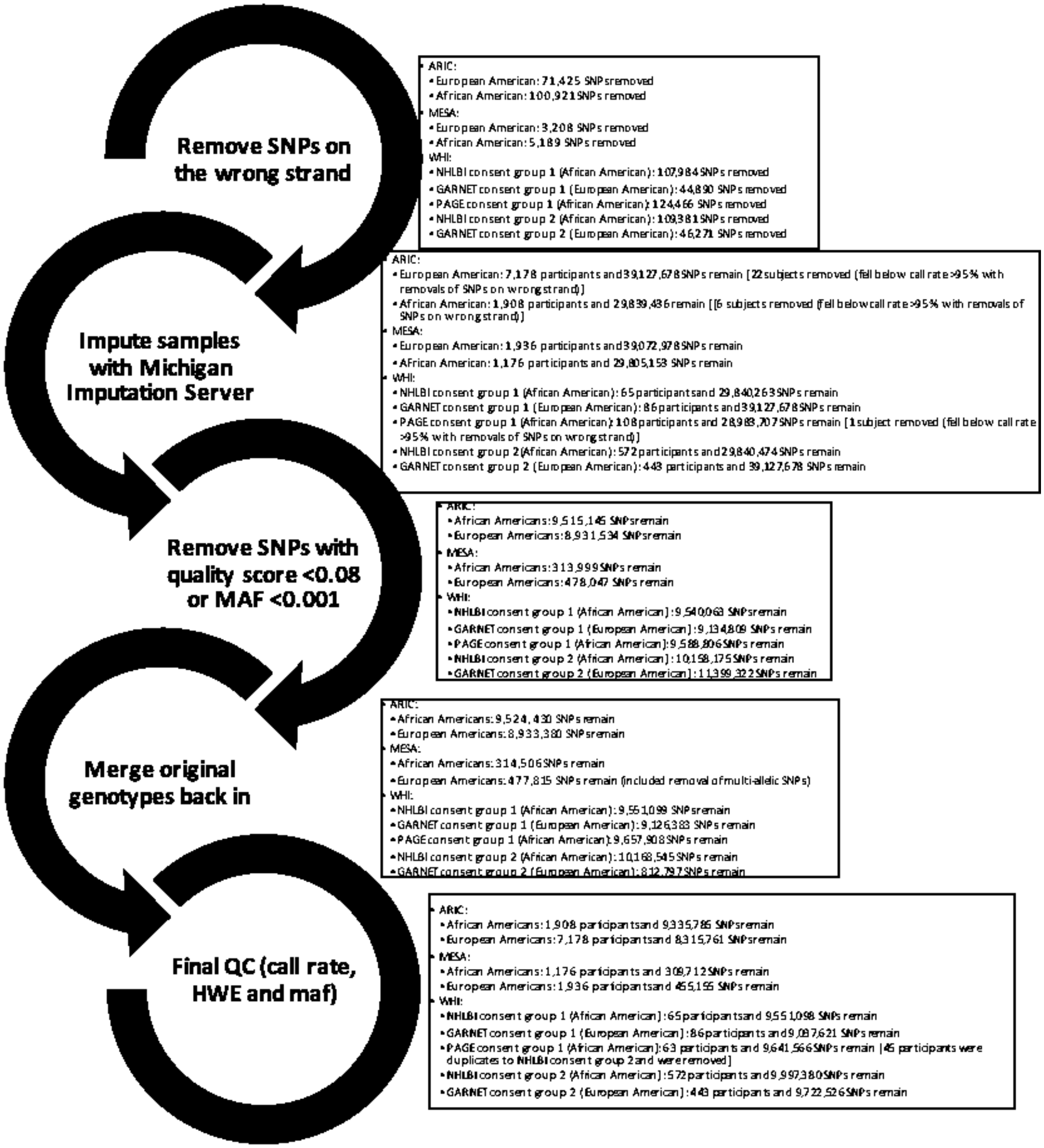


Abbreviations: SNP, single nucleotide polymorphism; HWE, Hardy Weinberg equilibrium; MAF, minor allele frequency; IBD, Identity-by-descent

Supplementary Figure 2 shows quality control steps and cutoffs used as well as corresponding sample sizes at each step starting at the imputation phase. Sample sizes reported here reflect entire eligible cohort, actual numbers used differ to maintain independence where necessary.

Supplementary Table 2: UV radiation values for ARIC*

| Field Center | | | | |
|---|---|---|---|---|
| Month of Visit | Wake Forest Baptist Medical Center, Winston-Salem, NC<br><br>Recruitment in Forsyth County, NC | University of Mississippi Medical Center, Jackson, MS<br><br>Recruitment in Jackson, MS | University of Minnesota, Minneapolis, MN<br><br>Recruitment in Northwestern Minneapolis, MN | Johns Hopkins University, Baltimore, MD<br><br>Recruitment in Washington County, MD |
| January | Month: December<br>Year: average 1994-2002<br>Location: Raleigh, NC | Month: December<br>Year: average 1994-2002<br>Location: Jackson, MS | Month: December<br>Year: average 1994-2002<br>Location: Minneapolis, MN | Month: December<br>Year: average 1994-2002<br>Location: Baltimore, MD and Pittsburgh, PA |
| February | Month: January<br>Year: average 1994-2002<br>Location: Raleigh, NC | Month: January<br>Year: average 1994-2002<br>Location: Jackson, MS | Month: January<br>Year: average 1994-2002<br>Location: Minneapolis, MN | Month: January<br>Year: average 1994-2002<br>Location: Baltimore, MD and Pittsburgh, PA |
| March | Month: February<br>Year: average 1994-2002<br>Location: Raleigh, NC | Month: February<br>Year: average 1994-2002<br>Location: Jackson, MS | Month: February<br>Year: average 1994-2002<br>Location: Minneapolis, MN | Month: February<br>Year: average 1994-2002<br>Location: Baltimore, MD and Pittsburgh, PA |
| April | Month: March<br>Year: average 1994-2002<br>Location: Raleigh, NC | Month: March<br>Year: average 1994-2002<br>Location: Jackson, MS | Month: March<br>Year: average 1994-2002<br>Location: Minneapolis, MN | Month: March<br>Year: average 1994-2002<br>Location: Baltimore, MD and Pittsburgh, PA |
| May | Month: April<br>Year: average 1994-2002<br>Location: Raleigh, NC | Month: April<br>Year: average 1994-2002<br>Location: Jackson, MS | Month: April<br>Year: average 1994-2002<br>Location: Minneapolis, MN | Month: April<br>Year: average 1994-2002<br>Location: Baltimore, MD and Pittsburgh, PA |
| June | Month: May<br>Year: average 1994-2002<br>Location: Raleigh, NC | Month: May<br>Year: average 1994-2002<br>Location: Jackson, MS | Month: May<br>Year: average 1994-2002<br>Location: R Minneapolis, MN | Month: May<br>Year: average 1994-2002<br>Location: Baltimore, MD and Pittsburgh, PA |
| July | Month: June<br>Year: average 1994-2002<br>Location: Raleigh, NC | Month: June<br>Year: average 1994-2002<br>Location: Jackson, MS | Month: June<br>Year: average 1994-2002<br>Location: Minneapolis, MN | Month: June<br>Year: average 1994-2002<br>Location: Baltimore, MD and Pittsburgh, PA |
| August | Month: July<br>Year: average 1994-2002<br>Location: Raleigh, NC | Month: July<br>Year: average 1994-2002<br>Location: Jackson, MS | Month: July<br>Year: average 1994-2002<br>Location: Minneapolis, MN | Month: July<br>Year: average 1994-2002<br>Location: Baltimore, MD and Pittsburgh, PA |
| September | Month: August<br>Year: average 1994-2002<br>Location: Raleigh, NC | Month: August<br>Year: average 1994-2002<br>Location: Jackson, MS | Month: August<br>Year: average 1994-2002<br>Location: Minneapolis, MN | Month: August<br>Year: average 1994-2002<br>Location: Baltimore, MD and Pittsburgh, PA |
| October | Month: September<br>Year: average 1994-2002<br>Location: Raleigh, NC | Month: September<br>Year: average 1994-2002<br>Location: Jackson, MS | Month: September<br>Year: average 1994-2002<br>Location: Minneapolis, MN | Month: September<br>Year: average 1994-2002<br>Location: Baltimore, MD and Pittsburgh, PA |
| November | Month: October<br>Year: average 1994-2002<br>Location: Raleigh, NC | Month: October<br>Year: average 1994-2002<br>Location: Jackson, MS | Month: October<br>Year: average 1994-2002<br>Location: Minneapolis, MN | Month: October<br>Year: average 1994-2002<br>Location: Baltimore, MD and Pittsburgh, PA |
| December | Month: November<br>Year: average 1994-2002<br>Location: Raleigh, NC | Month: November<br>Year: average 1994-2002<br>Location: Jackson, MS | Month: November<br>Year: average 1994-2002<br>Location: Minneapolis, MN | Month: November<br>Year: average 1994-2002<br>Location: Baltimore, MD and Pittsburgh, PA |

*All visits occurred between 1990 and 1993; the National Weather Service Climate Prediction Center database starts with June 1994, therefore, the average for years 1994-2002 (the years in which vitamin D was collected for this project) was used.

Supplemental Table 3: UV radiation values for MESA*

| | Site | | | | | |
|---|---|---|---|---|---|---|
| Month of visit | Wake Forest University, Winston-Salem, NC | Columbia University, New York, NY | Johns Hopkins University, Baltimore, MD | University of Minnesota, Minneapolis, MN | Northwestern University, Chicago, IL | University of California, Los Angeles, CA |
| January | Month: December Years: 2000-2002 Location: Raleigh, NC | Month: December Years: 2000-2002 Location: New York, NY | Month: December Years: 2000-2002 Location: Baltimore, MD | Month: December Years: 2000-2002 Location: Minneapolis, MN | Month: December Years: 2000-2002 Location: Chicago, IL | Month: December Years: 2000-2002 Location: Los Angeles, CA |
| February | Month: January Years: 2000-2002 Location: Raleigh, NC | Month: January Years: 2000-2002 Location: New York, NY | Month: January Years: 2000-2002 Location: Baltimore, MD | Month: January Years: 2000-2002 Location: Minneapolis, MN | Month: January Years: 2000-2002 Location: Chicago, IL | Month: January Years: 2000-2002 Location: Los Angeles, CA |
| March | Month: February Years: 2000-2002 Location: Raleigh, NC | Month: February Years: 2000-2002 Location: New York, NY | Month: February Years: 2000-2002 Location: Baltimore, MD | Month: February Years: 2000-2002 Location: Minneapolis, MN | Month: February Years: 2000-2002 Location: Chicago, IL | Month: February Years: 2000-2002 Location: Los Angeles, CA |
| April | Month: March Years: 2000-2002 Location: Raleigh, NC | Month: March Years: 2000-2002 Location: New York, NY | Month: March Years: 2000-2002 Location: Baltimore, MD | Month: March Years: 2000-2002 Location: Minneapolis, MN | Month: March Years: 2000-2002 Location: Chicago, IL | Month: March Years: 2000-2002 Location: Los Angeles, CA |
| May | Month: April Years: 2000-2002 Location: Raleigh, NC | Month: April Years: 2000-2002 Location: New York, NY | Month: April Years: 2000-2002 Location: Baltimore, MD | Month: April Years: 2000-2002 Location: Minneapolis, MN | Month: April Years: 2000-2002 Location: Chicago, IL | Month: April Years: 2000-2002 Location: Los Angeles, CA |
| June | Month: May Years: 2000-2002 Location: Raleigh, NC | Month: May Years: 2000-2002 Location: New York, NY | Month: May Years: 2000-2002 Location: Baltimore, MD | Month: May Years: 2000-2002 Location: Minneapolis, MN | Month: May Years: 2000-2002 Location: Chicago, IL | Month: May Years: 2000-2002 Location: Los Angeles, CA |
| July | Month: June Years: 2000-2002 Location: Raleigh, NC | Month: June Years: 2000-2002 Location: New York, NY | Month: June Years: 2000-2002 Location: Baltimore, MD | Month: June Years: 2000-2002 Location: Minneapolis, MN | Month: June Years: 2000-2002 Location: Chicago, IL | Month: June Years: 2000-2002 Location: Los Angeles, CA |
| August | Month: July Years: 2000-2002 Location: Raleigh, NC | Month: July Years: 2000-2002 Location: New York, NY | Month: July Years: 2000-2002 Location: Baltimore, MD | Month: July Years: 2000-2002 Location: Minneapolis, MN | Month: July Years: 2000-2002 Location: Chicago, IL | Month: July Years: 2000-2002 Location: Los Angeles, CA |
| September | Month: August Years: 2000-2002 Location: Raleigh, NC | Month: August Years: 2000-2002 Location: New York, NY | Month: August Years: 2000-2002 Location: Baltimore, MD | Month: August Years: 2000-2002 Location: Minneapolis, MN | Month: August Years: 2000-2002 Location: Chicago, IL | Month: August Years: 2000-2002 Location: Los Angeles, CA |
| October | Month: September Years: 2000-2002 Location: Raleigh, NC | Month: September Years: 2000-2002 Location: New York, NY | Month: September Years: 2000-2002 Location: Baltimore, MD | Month: September Years: 2000-2002 Location: Minneapolis, MN | Month: September Years: 2000-2002 Location: Chicago, IL | Month: September Years: 2000-2002 Location: Los Angeles, CA |
| November | Month: October Years: 2000-2002 Location: Raleigh, NC | Month: October Years: 2000-2002 Location: New York, NY | Month: October Years: 2000-2002 Location: Baltimore, MD | Month: October Years: 2000-2002 Location: Minneapolis, MN | Month: October Years: 2000-2002 Location: Chicago, IL | Month: October Years: 2000-2002 Location: Los Angeles, CA |
| December | Month: November Years: 2000-2002 Location: Raleigh, NC | Month: November Years: 2000-2002 Location: New York, NY | Month: November Years: 2000-2002 Location: Baltimore, MD | Month: November Years: 2000-2002 Location: Minneapolis, MN | Month: November Years: 2000-2002 Location: Chicago, IL | Month: November Years: 2000-2002 Location: Los Angeles, CA |

*Month, but not specific year variables available for all participants, however, all visits occurred between 2000-2002, therefore, monthly average for years 2000-2002 was used.

Supplemental Table 4: UV radiation values for WHI*

| | Location | | | | | | | | |
|---|---|---|---|---|---|---|---|---|---|
| Month of visit | Northeast (35-40 degrees N) | Northeast (>40 degrees N) | South (<35 degrees N) | South (35-40 degrees N) | Midwest (35-40 degrees N) | Midwest (>40 degrees N) | West (<35 degrees N) | West (35-40 degrees N) | West (>40 degrees N) |
| January | Month: December Locations: Raleigh, NC; Charleston, SC; Washington, DC | Month: December Locations: New York, NY; Buffalo, NY; Burlington, VT; Boston, MA; Portland, ME | Month: December Locations: Los Angeles, CA; Phoenix, AZ; Houston, TX; Atlanta, GA; Jacksonville, FL; Miami, FL | Month: December Locations: San Francisco, CA; Memphis, TN | Month: December Locations: St. Louis, MO; Omaha, NE; Sioux Falls, SD | Month: December Locations: Milwaukee, WI; Minneapolis, MN; Chicago, IL; Des Moines, IA; Bismarck, ND | Month: December Locations: Phoenix, AZ; Los Angeles, CA; Albuquerque, NM | Month: December Locations: San Francisco, CA; Denver, CO; Salt Lake City, UT; Las Vegas, NV | Month: December Locations: Portland, OR; Seattle, WA; Billing, MT; Boise, ID |
| February | Month: January Locations: NC; Charleston, SC; Washington, DC | Month: January Locations: New York, NY; Buffalo, NY; Burlington, VT; Boston, MA; Portland, ME | Month: January Locations: Los Angeles, CA; Phoenix, AZ; Houston, TX; Atlanta, GA; Jacksonville, FL; Miami, FL | Month: January Locations: San Francisco, CA; Memphis, TN | Month: January Locations: St. Louis, MO; Omaha, NE; Sioux Falls, SD | Month: January Locations: Milwaukee, WI; Minneapolis, MN; Chicago, IL; Des Moines, IA; Bismarck, ND | Month: January Locations: Phoenix, AZ; Los Angeles, CA; Albuquerque, NM | Month: January Locations: San Francisco, CA; Denver, CO; Salt Lake City, UT; Las Vegas, NV | Month: January Locations: Portland, OR; Seattle, WA; Billing, MT; Boise, ID |
| March | Month: February Locations: NC; Charleston, SC; Washington, DC | Month: February Locations: New York, NY; Buffalo, NY; Burlington, VT; Boston, MA; Portland, ME | Month: February Locations: Los Angeles, CA; Phoenix, AZ; Houston, TX; Atlanta, GA; Jacksonville, FL; Miami, FL | Month: February Locations: San Francisco, CA; Memphis, TN | Month: February Locations: St. Louis, MO; Omaha, NE; Sioux Falls, SD | Month: February Locations: Milwaukee, WI; Minneapolis, MN; Chicago, IL; Des Moines, IA; Bismarck, ND | Month: February Locations: Phoenix, AZ; Los Angeles, CA; Albuquerque, NM | Month: February Locations: San Francisco, CA; Denver, CO; Salt Lake City, UT; Las Vegas, NV | Month: February Locations: Portland, OR; Seattle, WA; Billing, MT; Boise, ID |
| April | Month: March Locations: NC; Charleston, SC; Washington, DC | Month: March Locations: New York, NY; Buffalo, NY; Burlington, VT; Boston, MA; Portland, ME | Month: March Locations: Los Angeles, CA; Phoenix, AZ; Houston, TX; Atlanta, GA; Jacksonville, FL; Miami, FL | Month: March Locations: San Francisco, CA; Memphis, TN | Month: March Locations: St. Louis, MO; Omaha, NE; Sioux Falls, SD | Month: March Locations: Milwaukee, WI; Minneapolis, MN; Chicago, IL; Des Moines, IA; Bismarck, ND | Month: March Locations: Phoenix, AZ; Los Angeles, CA; Albuquerque, NM | Month: March Locations: San Francisco, CA; Denver, CO; Salt Lake City, UT; Las Vegas, NV | Month: March Locations: Portland, OR; Seattle, WA; Billing, MT; Boise, ID |
| May | Month: April Locations: NC; Charleston, SC; Washington, DC | Month: April Locations: New York, NY; Buffalo, NY; Burlington, VT; Boston, MA; Portland, ME | Month: April Locations: Los Angeles, CA; Phoenix, AZ; Houston, TX; Atlanta, GA; Jacksonville, FL; Miami, FL | Month: April Locations: San Francisco, CA; Memphis, TN | Month: April Locations: St. Louis, MO; Omaha, NE; Sioux Falls, SD | Month: April Locations: Milwaukee, WI; Minneapolis, MN; Chicago, IL; Des Moines, IA; Bismarck, ND | Month: April Locations: Phoenix, AZ; Los Angeles, CA; Albuquerque, NM | Month: April Locations: San Francisco, CA; Denver, CO; Salt Lake City, UT; Las Vegas, NV | Month: April Locations: Portland, OR; Seattle, WA; Billing, MT; Boise, ID |
| June | Month: May Locations: NC; Charleston, SC; Washington, DC | Month: May Locations: New York, NY; Buffalo, NY; Burlington, VT; Boston, | Month: May Locations: Los Angeles, CA; Phoenix, AZ; Houston, TX; Atlanta, GA; | Month: May Locations: San Francisco, CA; Memphis, TN | Month: May Locations: St. Louis, MO; Omaha, NE; Sioux Falls, SD | Month: May Locations: Milwaukee, WI; Minneapolis, MN; Chicago, IL; Des | Month: May Locations: Phoenix, AZ; Los Angeles, CA; Albuquerque, NM | Month: May Locations: San Francisco, CA; Denver, CO; Salt Lake City, | Month: May Locations: Portland, OR; Seattle, WA; Billing, MT; Boise, ID |

| | | | | | | | | | |
|---|---|---|---|---|---|---|---|---|---|
| | | MA; Portland, ME | Jacksonville, FL; Miami, FL | | | Moines, IA; Bismarck, ND | | UT; Las Vegas, NV | |
| July | Month: June Locations: NC; Charleston, SC; Washington, DC | Month: June Locations: New York, NY; Buffalo, NY; Burlington, VT; Boston, MA; Portland, ME | Month: June Locations: Los Angeles, CA; Phoenix, AZ; Houston, TX; Atlanta, GA; Jacksonville, FL; Miami, FL | Month: June Locations: San Francisco, CA; Memphis, TN | Month: June Locations: St. Louis, MO; Omaha, NE; Sioux Falls, SD | Month: June Locations: Milwaukee, WI; Minneapolis, MN; Chicago, IL; Des Moines, IA; Bismarck, ND | Month: June Locations: Phoenix, AZ; Los Angeles, CA; Albuquerque, NM | Month: June Locations: San Francisco, CA; Denver, CO; Salt Lake City, UT; Las Vegas, NV | Month: June Locations: Portland, OR; Seattle, WA; Billing, MT; Boise, ID |
| August | Month: July Locations: NC; Charleston, SC; Washington, DC | Month: July Locations: New York, NY; Buffalo, NY; Burlington, VT; Boston, MA; Portland, ME | Month: July Locations: Los Angeles, CA; Phoenix, AZ; Houston, TX; Atlanta, GA; Jacksonville, FL; Miami, FL | Month: July Locations: San Francisco, CA; Memphis, TN | Month: July Locations: St. Louis, MO; Omaha, NE; Sioux Falls, SD | Month: July Locations: Milwaukee, WI; Minneapolis, MN; Chicago, IL; Des Moines, IA; Bismarck, ND | Month: July Locations: Phoenix, AZ; Los Angeles, CA; Albuquerque, NM | Month: July Locations: San Francisco, CA; Denver, CO; Salt Lake City, UT; Las Vegas, NV | Month: July Locations: Portland, OR; Seattle, WA; Billing, MT; Boise, ID |
| September | Month: August Locations: NC; Charleston, SC; Washington, DC | Month: August Locations: New York, NY; Buffalo, NY; Burlington, VT; Boston, MA; Portland, ME | Month: August Locations: Los Angeles, CA; Phoenix, AZ; Houston, TX; Atlanta, GA; Jacksonville, FL; Miami, FL | Month: August Locations: San Francisco, CA; Memphis, TN | Month: August Locations: St. Louis, MO; Omaha, NE; Sioux Falls, SD | Month: August Locations: Milwaukee, WI; Minneapolis, MN; Chicago, IL; Des Moines, IA; Bismarck, ND | Month: August Locations: Phoenix, AZ; Los Angeles, CA; Albuquerque, NM | Month: August Locations: San Francisco, CA; Denver, CO; Salt Lake City, UT; Las Vegas, NV | Month: August Locations: Portland, OR; Seattle, WA; Billing, MT; Boise, ID |
| October | Month: September Locations: NC; Charleston, SC; Washington, DC | Month: September Locations: New York, NY; Buffalo, NY; Burlington, VT; Boston, MA; Portland, ME | Month: September Locations: Los Angeles, CA; Phoenix, AZ; Houston, TX; Atlanta, GA; Jacksonville, FL; Miami, FL | Month: September Locations: San Francisco, CA; Memphis, TN | Month: September Locations: St. Louis, MO; Omaha, NE; Sioux Falls, SD | Month: September Locations: Milwaukee, WI; Minneapolis, MN; Chicago, IL; Des Moines, IA; Bismarck, ND | Month: September Locations: Phoenix, AZ; Los Angeles, CA; Albuquerque, NM | Month: September Locations: San Francisco, CA; Denver, CO; Salt Lake City, UT; Las Vegas, NV | Month: September Locations: Portland, OR; Seattle, WA; Billing, MT; Boise, ID |
| November | Month: October Locations: NC; Charleston, SC; Washington, DC | Month: October Locations: New York, NY; Buffalo, NY; Burlington, VT; Boston, MA; Portland, ME | Month: October Locations: Los Angeles, CA; Phoenix, AZ; Houston, TX; Atlanta, GA; Jacksonville, FL; Miami, FL | Month: October Locations: San Francisco, CA; Memphis, TN | Month: October Locations: St. Louis, MO; Omaha, NE; Sioux Falls, SD | Month: October Locations: Milwaukee, WI; Minneapolis, MN; Chicago, IL; Des Moines, IA; Bismarck, ND | Month: October Locations: Phoenix, AZ; Los Angeles, CA; Albuquerque, NM | Month: October Locations: San Francisco, CA; Denver, CO; Salt Lake City, UT; Las Vegas, NV | Month: October Locations: Portland, OR; Seattle, WA; Billing, MT; Boise, ID |
| December | Month: November Locations: NC; Charleston, SC; Washington, DC | Month: November Locations: New York, NY; Buffalo, NY; Burlington, VT; Boston, MA; Portland, ME | Month: November Locations: Los Angeles, CA; Phoenix, AZ; Houston, TX; Atlanta, GA; Jacksonville, FL; Miami, FL | Month: November Locations: San Francisco, CA; Memphis, TN | Month: November Locations: St. Louis, MO; Omaha, NE; Sioux Falls, SD | Month: November Locations: Milwaukee, WI; Minneapolis, MN; Chicago, IL; Des Moines, IA; Bismarck, ND | Month: November Locations: Phoenix, AZ; Los Angeles, CA; Albuquerque, NM | Month: November Locations: San Francisco, CA; Denver, CO; Salt Lake City, UT; Las Vegas, NV | Month: November Locations: Portland, OR; Seattle, WA; Billing, MT; Boise, ID |

*Blood draws for WHI were done in the years 1993-1999, the years used for UV radiation value match the year of visit, with the exception of 1993 and 1994 (before the National Weather Service Climate Prediction Center database had started documenting data,. Those with visits in 1993 or 1994 are given the corresponding monthly average for the average of years 1995-2002.

Supplemental Table 5: Available UV radiation value descriptive statistics for ARIC (average from 1994-2002)

| Field Center | | | | |
|---|---|---|---|---|
| Month of blood draw | Wake Forest Baptist Medical Center, Winston-Salem, NC | University of Mississippi Medical Center, Jackson, MS | University of Minnesota, Minneapolis, MN | Johns Hopkins University, Baltimore, MD |
| January | 2.1 | 2.7 | 0.9 | 1.4 |
| February | 2.5 | 3.2 | 1.1 | 1.8 |
| March | 3.7 | 4.7 | 1.9 | 2.8 |
| April | 5.7 | 6.8 | 3.4 | 4.5 |
| May | 7.1 | 8.2 | 4.8 | 6.1 |
| June | 8.1 | 8.9 | 6.2 | 7.3 |
| July | 8.7 | 9.1 | 7.5 | 8.3 |
| August | 8.9 | 9.5 | 7.8 | 8.4 |
| September | 8.3 | 9.1 | 6.7 | 7.5 |
| October | 6.6 | 7.7 | 4.7 | 5.7 |
| November | 4.5 | 5.5 | 2.5 | 3.6 |
| December | 2.8 | 3.5 | 1.3 | 2.0 |

Supplemental Table 6: Available UV radiation value descriptive statistics for MESA (average from 2000-2002)

| Site | | | | | | |
|---|---|---|---|---|---|---|
| Month of visit | Wake Forest University, Winston-Salem, NC | Columbia University, New York, NY | Johns Hopkins University, Baltimore, MD | University of Minnesota, Minneapolis, MN | Northwestern University, Chicago, IL | University of California, Los Angeles, CA |
| January | 1.9 | 1.1 | 1.5 | 0.7 | 0.9 | 2.3 |
| February | 2.5 | 1.6 | 1.9 | 1.1 | 1.5 | 2.8 |
| March | 3.8 | 2.6 | 2.9 | 2.0 | 2.5 | 4.1 |
| April | 5.9 | 4.4 | 4.8 | 3.3 | 4.0 | 5.9 |
| May | 6.9 | 5.4 | 6.3 | 4.6 | 5.3 | 7.3 |
| June | 7.7 | 6.4 | 7.1 | 5.6 | 6.2 | 8.2 |
| July | 8.4 | 7.6 | 8.2 | 6.9 | 7.5 | 8.9 |
| August | 8.1 | 7.2 | 8.0 | 7.3 | 7.7 | 9.1 |
| September | 7.6 | 6.5 | 7.3 | 6.1 | 6.7 | 8.9 |
| October | 6.0 | 5.0 | 5.7 | 4.3 | 4.9 | 7.3 |
| November | 4.3 | 3.0 | 3.7 | 2.1 | 2.6 | 4.5 |
| December | 2.5 | 1.6 | 2.0 | 1.1 | 2.4 | 2.9 |

Supplemental Table 7: Available UV radiation value descriptive statistics for WHI^

| | Location | | | | | | | | |
|---|---|---|---|---|---|---|---|---|---|
| Month of visit | Northeast (35-40 degrees N) | Northeast (>40 degrees N) | South (<35 degrees N) | South (35-40 degrees N) | Midwest (35-40 degrees N) | Midwest (>40 degrees N) | West (<35 degrees N) | West (35-40 degrees N) | West (>40 degrees N) |
| January | 1993-4*:1.8 | 1993-4*: 1.1<br>1996: 1.2 | 1993-4*: 2.6<br>1999: 2.2 | 1993-4*: 2.0<br>1996: 2.2<br>1999: 1.5 | 1993-4*: 1.3<br>1999: 0.9 | 1993-4*: 1.0<br>1996: 1.1<br>1999: 0.7 | 1993-4*: 2.5<br>1999: 2.2 | 1993-4*: 1.8<br>1999: 1.4 | 1993-4*: 0.8<br>1996: 0.9 |
| February | 1993-4*: 2.2<br>1999: 2.1 | 1993-4*: 1.3<br>1999: 1.3 | 1993-4*: 3.1<br>1996: 3.3 | 1993-4*: 2.4<br>1996: 2.5 | 1993-4*: 1.6 | 1993-4*: 1.3 | 1993-4*: 2.9<br>1999: 2.9 | 1993-4*: 2.1<br>1999: 2.1 | 1993-4*: 1.1<br>1996: 1.1 |
| March | 1993-4*: 3.3 | 1993-4*: 2.2<br>1999: 2.2 | 1993-4*:4.5<br>1996: 4.5<br>1999: 4.5 | 1993-4*: 3.6<br>1996: 3.6 | 1993-4*: 2.6<br>1999: 2.5 | 1993-4*: 2.1<br>1999: 2.1 | 1993-4*: 4.2 | 1993-4*:3.2<br>1996: 3.3<br>1999: 3.3 | 1993-4*: 2.0 |
| April | 1993-4*: 5.1 | 1993-4*: 3.7<br>1996: 3.8 | 1993-4*: 6.4 | 1993-4*: 5.5<br>1996: 5.4 | 1993-4*: 4.3<br>1996: 4.2 | 1993-4*: 3.7<br>1996: 3.8<br>1999: 3.6 | 1993-4*: 6.1<br>1999: 5.9 | 1993-4*: 5.1 | 1993-4*: 3.4 |
| May | 1993-4*: 6.6 | 1993-4*: 5.2<br>1999: 4.2 | 1993-4*: 7.7<br>1996: 8.3<br>1999: 6.4 | 1993-4*: 7.0 | 1993-4*: 5.7 | 1993-4*: 5.2<br>1996: 5.6<br>1999: 4.4 | 1993-4*: 7.7 | 1993-4*: 6.6<br>1999: 5.4 | 1993-4*: 4.8 |
| June | 1993-4*: 7.7 | 1993-4*: 6.6<br>1999: 7.1 | 1993-4*: 8.5<br>1999: 7.4 | 1993-4*: 8.0<br>1996: 9.1 | 1993-4*: 7.2 | 1993-4*: 6.6<br>1996: 7.5<br>1999: 5.1 | 1993-4*: 8.7 | 1993-4*: 8.1<br>1999: 7.2 | 1993-4*: 6.3 |
| July | 1993-4*: 8.5 | 1993-4*: 7.8<br>1999: 6.6 | 1993-4*: 9.0<br>1996: 10.0 | 1993-4*: 8.8<br>1996: 9.5<br>1999: 7.5 | 1993-4*: 8.3 | 1993-4*: 7.8 | 1993-4*: 8.9 | 1993-4*: 8.8<br>1996: 9.9 | 1993-4*: 7.3<br>1996: 8.2 |
| August | 1993-4*: 8.7 | 1993-4*: 7.7<br>1996: 8.4<br>1999: 6.8 | 1993-4*: 9.1<br>1996: 9.9<br>1999: 8.0 | 1993-4*: 9.1<br>1999: 8.0 | 1993-4*: 8.6<br>1996: 9.1<br>1999:7 .5 | 1993-4*: 8.1<br>1996: 8.5 | 1993-4*: 9.2 | 1993-4*: 9.1 | 1993-4*: 7.7 |
| September | 1993-4*: 7.9<br>1996: 8.7 | 1993-4*: 6.7<br>1999: 5.3 | 1993-4*: 8.8<br>1999: 7.8 | 1993-4*:8.5<br>1996: 9.2 | 1993-4*: 7.6<br>1996: 8.3 | 1993-4*: 7.0<br>1996: 7.7<br>1999: 6.0 | 1993-4*: 9.2 | 1993-4*: 8.6<br>1999: 7.9 | 1993-4*: 6.7 |
| October | 1993-4*: 6.2<br>1999: 5.0 | 1993-4*: 5.0<br>1996: 5.5 | 1993-4*: 7.4<br>1999: 6.5 | 1993-4*: 6.7 | 1993-4*: 5.6<br>1996: 5.9<br>1999: 4.7 | 1993-4*: 5.0<br>1996: 5.3<br>1999: 4.0 | 1993-4*: 7.7<br>1996: 7.9<br>1999: 7.1 | 1993-4*: 6.7<br>1999: 6.0 | 1993-4*: 4.7<br>1996: 4.9 |
| November | 1993-4*: 4.1 | 1993-4*: 2.9 | 1993-4*: 5.2 | 1993-4*: 4.4 | 1993-4*: 3.3 | 1993-4*: 2.8 | 1993-4*: 5.1 | 1993-4*: 4.1 | 1993-4*: 2.6 |

| | | | | | | | | | |
|---|---|---|---|---|---|---|---|---|---|
| | | 1996: 3.1<br>1999: 2.3 | 1996: 5.5<br>1999: 4.5 | 1996: 4.7<br>1999: 4.0 | | 1996: 3.1 | 1996: 5.2 | | |
| December | 1993-4*: 2.4 | 1993-4*: 1.6<br>1996: 1.7<br>1999: 1.2 | 1993-4*: 3.4<br>1996: 3.8<br>1999: 2.9 | 1993-4*: 2.7 | 1993-4*: 1.9<br>1996: 2.1 | 1993-4*: 1.5<br>1996: 1.7<br>1999: 1.3 | N/A | 1993-4*: 2.4 | 1993-4*: 1.3 |

^Only values for location, month and year combos with observations in the final dataset are shown.

*Average of 1995-2002 (all data collected, since database started in late 1994)

Supplemental Table 8: Performance of models in determining optimal p-value cutoff

| **Ancestry** | **Model** | **Cut-off** | **PGS $R^2$ (model $R^2$)** | **p-value** | **# SNPs in PGS** |
|---|---|---|---|---|---|
| **European (n=1,000)** | **No covariates** | 0.00025 | 0.015 (0.015) | 0.00009 | 251 |
| | **Age + sex** | 0.00025 | 0.016 (0.052) | 0.00005 | 251 |
| | **Age + sex +BMI** | 0.00035 | 0.016 (0.065) | 0.00005 | 341 |
| | **Age + sex + UV** | 0.00035 | 0.014 (0.106) | 0.000001 | 341 |
| | **Age + sex + BMI +UV[a]** | 0.00035 | 0.014 (0.129) | 0.00008 | 341 |
| | **Age + sex + BMI +UV + intake** | 0.00035 | 0.013 (0.137) | 0.00007 | 341 |
| **African (n=57)** | **No covariates** | 0.01265 | 0.081 (0.081) | 0.024 | 32,269 |
| | **Age + sex** | 0.01265 | 0.107 (0.179) | 0.008 | 32,269 |
| | **Age + sex +BMI** | 0.01265 | 0.08 (0.162) | 0.03 | 32,269 |
| | **Age + sex + UV** | 0.01265 | 0.105 (0.273) | 0.006 | 32,269 |
| | **Age + sex + BMI +UV[a]** | 0.01265 | 0.044 (0.37) | 0.06 | 32,269 |
| | **Age + sex + BMI +UV + intake** | 0.0072 | 0.011 (0.33) | 0.47 | 19,261 |

Abbreviations: BMI, body mass index; UV, ultra-violet; PGS, polygenic score; SNP, single nucleotide polymorphism

[a]Optimal model utilized in analyses

Supplemental Table 9: Performance of optimal PGS in each ancestry utilizing LD cutoff of 0.2

| **Ancestry** | **p-value cut-off** | **# SNPs** | **PGS $R^2$ (model $R^2$)** | **p-value[a]** |
|---|---|---|---|---|
| **European** | 0.1142 | 44,883 | 0.010 (0.13) | $5.8x10^{-4}$ |
| **African** | 0.01265 | 25,750 | 0.025 (0.45) | 0.14 |

Abbreviations: LD, linkage disequilibrium; SNPs, single nucleotide polymorphism; PGS, polygenic score

[a]p-value for association between PGS and log[25(OH)D]

Supplemental Table 10: Performance of unrestricted PGS in those of African ancestry

| **Target ancestry** | **Base ancestry** | **p-value cut-off** | **PGS $R^2$ (model $R^2$)** | **p-value[a]** |
|---|---|---|---|---|
| African | African | 1.0 | 0.0031 (0.0970) | 0.05 |
| | European | 1.0 | 0.0014 (0.0953) | 0.19 |

Abbreviations: SNPs, single nucleotide polymorphism; PGS, polygenic score

[a]p-value for association between PGS and log[25(OH)D]

Supplemental Figure 3: Visual representation of the association between PGS quintile and normalized vitamin D concentrations in those of African ancestry (n=1,042)

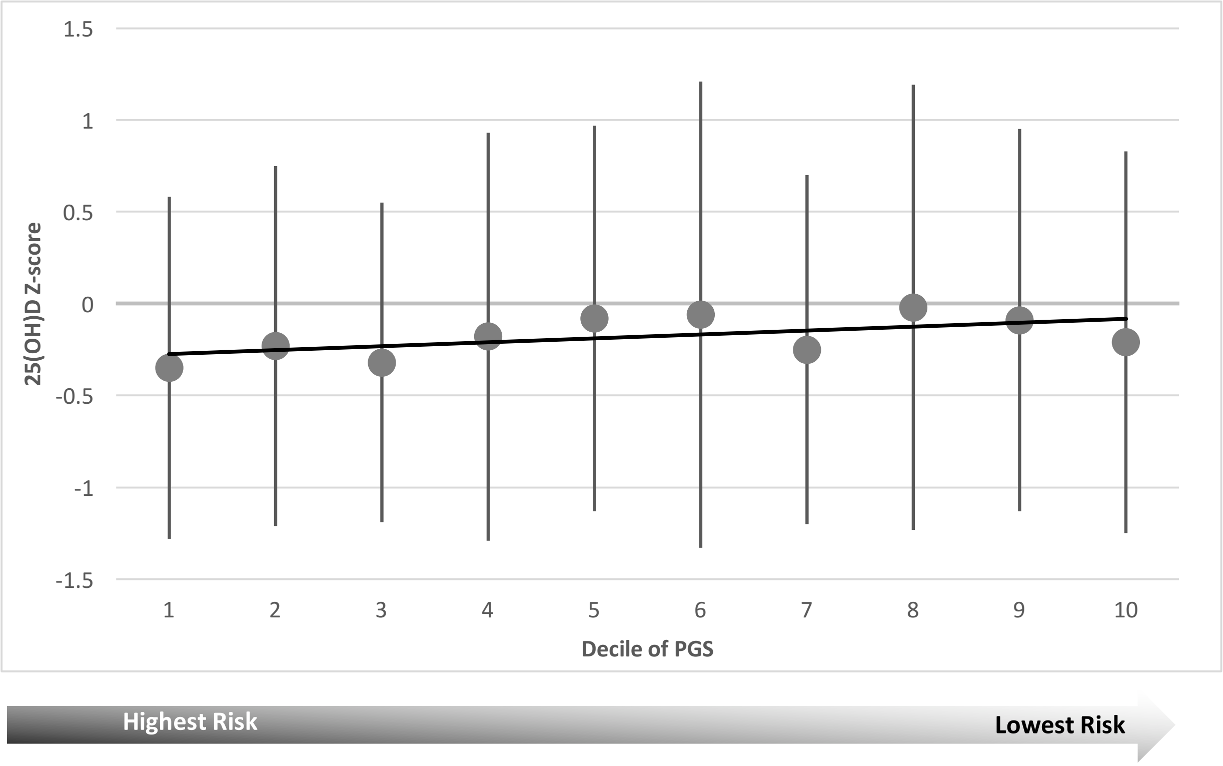


Supplemental Figure 3 is a visual representation of the association between polygenic score (PGS) quintile and normalized vitamin D concentrations in those of African ancestry (n=1,042). The x-axis is the PGS quintile, where lower quintile means more risk of low vitamin D concentrations. The y-axis is vitamin D concentrations (normalized for comparison between cohorts). As the PGS decreases (i.e. higher genetic risk) 25(OH)D concentrations decrease. Moving from the lowest risk to the highest risk decile decreases vitamin D concentrations by 1.9 ng/ml (p=0.37).

Supplemental Figure 4: Overall SNP heritability and replicated GWAS SNP heritability in those of African ancestry

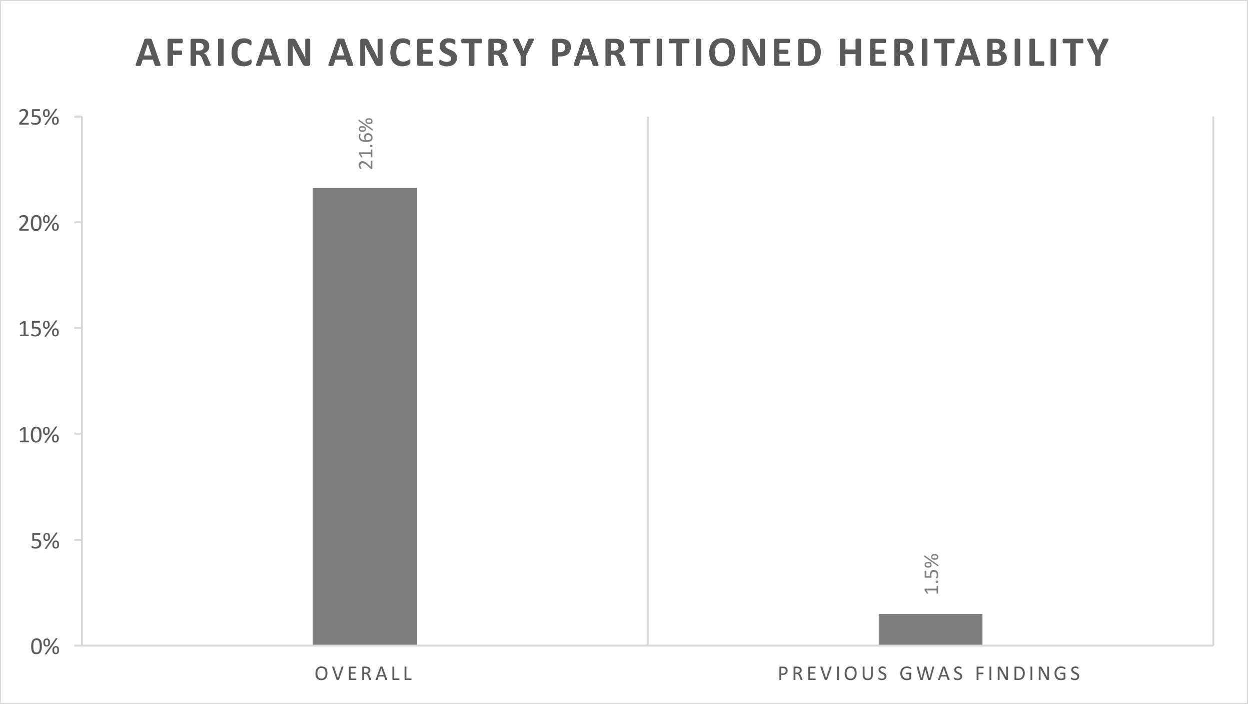


Supplemental Figure 4 shows overall single nucleotide polymorphism (SNP) heritability and replicated genome-wide association study (GWAS) SNP heritability for those of African ancestry. Sample size did not allow for calculation of the heritability of the PGS. Previous replicated GWAS findings, leave much of the heritability unexplained.
